# *Staphylococcal* secreted cytotoxins are competition sensing signals for *Pseudomonas aeruginosa*

**DOI:** 10.1101/2023.01.29.526047

**Authors:** Grace Z. Wang, Elizabeth A. Warren, Allison L. Haas, Andrea Sánchez Peña, Megan R. Kiedrowski, Brett Lomenick, Tsui-Fen Chou, Jennifer M. Bomberger, David A. Tirrell, Dominique H. Limoli

## Abstract

Coinfection with two notorious opportunistic pathogens, the Gram-negative *Pseudomonas aeruginosa* and Gram-positive *Staphylococcus aureus*, dominates chronic pulmonary infections. While coinfection is associated with poor patient outcomes, the interspecies interactions responsible for such decline remain unknown. Here, we dissected molecular mechanisms of interspecies sensing between *P. aeruginosa* and *S. aureus*. We discovered that *P. aeruginosa* senses *S. aureus* secreted peptides and, counterintuitively, moves towards these toxins. *P. aeruginosa* tolerates such a strategy through “competition sensing”, whereby it preempts imminent danger/competition by arming cells with type six secretion (T6S) and iron acquisition systems. Intriguingly, while T6S is predominantly described as weaponry targeting Gram-negative and eukaryotic cells, we find that T6S is essential for full *P. aeruginosa* competition with *S. aureus*, a previously undescribed role for T6S. Importantly, competition sensing was activated during coinfection of bronchial epithelia, including T6S islands targeting human cells. This study reveals critical insight into both interspecies competition and how antagonism may cause collateral damage to the host environment.

## INTRODUCTION

The future of microbiome research lies in our ability to manipulate polymicrobial interactions toward improved human health outcomes, which requires a fundamental molecular understanding of how microbial species sense and respond to ecological competition. Chronic respiratory infections in people with cystic fibrosis (CF) consist of diverse and heterogeneous microbial communities^1^. Nonetheless, *Pseudomonas aeruginosa* and *Staphylococcus aureus* are the most prevalent^2,3^. Critically, coinfection with these pathogens correlates with worsened clinical outcome and altered antibiotic efficacy^2-4^, urging the need for molecular dissection of their interspecies crosstalk.

We previously reported that *P. aeruginosa* is attracted to *S. aureus* resulting in invasion of *S. aureus* colonies^5^; however, what, if any, selective benefit *P. aeruginosa* achieves by adopting this behavior remains unknown. A potential role for such a strategy may be to bridge cellular distances for contact-dependent mechanisms of antagonism. The type six secretion system (T6SS), widely found in Gram-negative bacteria, such as *P. aeruginosa*, equips cells with a versatile nanomachinery that functions as an interspecies weapon capable of targeting both eukaryotic and prokaryotic cells^6-8^. *P. aeruginosa* typically maintains low basal T6SS activity but is capable of rapid reciprocal firing following T6SS attack by other Gram-negative species^9,10^. However, whether an analogous response may occur in response to Gram-positive competitors lacking T6SS remains unknown. A greater fundamental understanding of interspecies pathogen sensing and resulting competition, particularly between Gram negative and positive pathogens common during coinfection, is necessary to develop interventions directed at interspecies interactions.

Here, we report the discovery that *P. aeruginosa* rapidly activates T6SS after an encounter with the Gram-positive pathogen *S. aureus*. We present a “competition sensing” model uncovered by a combination of genetics, microscopy and multi-omics approaches whereby secreted *Staphylococcal* peptides are key interspecies signals that trigger *P. aeruginosa* antagonism. *P. aeruginosa* was found to sense *S. aureus* via secreted peptides at a distance, subsequently increasing directional motility and activating T6SS antagonism. Surprisingly, such activation allowed for T6SS-dependent competition with *S. aureus*, extending the functional role of T6SS to not only competition between Gram-negatives, but also between Gram-negative and positive bacteria. Furthermore, we examined coinfection on fully differentiated CF-derived bronchial epithelia, the gold standard model of *in vivo* CF airway infection, and found *P. aeruginosa* T6SS was activated, including host-targeting T6SS islands. Overall, these results broaden our mechanistic understanding of interspecies antagonism between distantly related species, reveal interspecies pathways that might be targeted therapeutically, and lend insight into the mechanism of increased patient decline during coinfection with *P. aeruginosa* and *S. aureus*.

## RESULTS

### PSMɑ peptides are necessary and sufficient for *P. aeruginosa* attraction toward *S. aureus*

We previously reported that *P. aeruginosa* travels up a gradient of *S. aureus* secreted factors using type-IV pilus (TFP)-dependent motility^5^. The *S. aureus* attractants identified are secreted *S. aureus* peptides, referred to as phenol soluble modulin (PSMs). *S. aureus* produces five alpha peptides: PSMɑ1-4 and PSMδ (δ toxin) and two β-peptides: PSMβ1 and 2 (**Supplementary Fig. 1**). Here, we first asked if *P. aeruginosa* possesses specificity in attraction towards individual peptides in a macroscopic TFP chemotaxis assay (**Fig.1a**). PSMɑ peptides were examined for initial characterization given that the ɑ peptides have known roles in neutrophil chemoattraction^11^ and cytotoxicity to mammalian host cells^12^. *P. aeruginosa* traveled further towards an increasing gradient of WT *S. aureus* supernatant (**Fig. 1b**), whereas directional motility towards supernatant derived from a double *psm*ɑ1-4 and *psm*δ mutant (Δ*psm*ɑ1-4 δATG-ATT, **Supplementary Fig. 1**) was eliminated, suggesting that at least one ɑ-peptide is necessary for attracting *P. aeruginosa* (**Fig. 1b, c**). The magnitude of attraction towards Δ*psm*ɑ1-4 was between WT *S. aureus* and the double Δ*psm*ɑ1-4 δATG-ATT mutants, suggesting PSMδ, along with the other ɑ peptides, is necessary for *P. aeruginosa* directional motility. We then determined if PSMs are sufficient to attract *P. aeruginosa* and the specificity of individual PSM peptide’s contribution. Pure synthetic PSMɑ3 and δ-toxin strongly attracted *P. aeruginosa* in a dose-dependent manner (**Fig. 1d**). These data demonstrate that PSMδ and PSMɑ3 are necessary and sufficient for *S. aureus* to attract *P. aeruginosa*.

**Figure 1.**
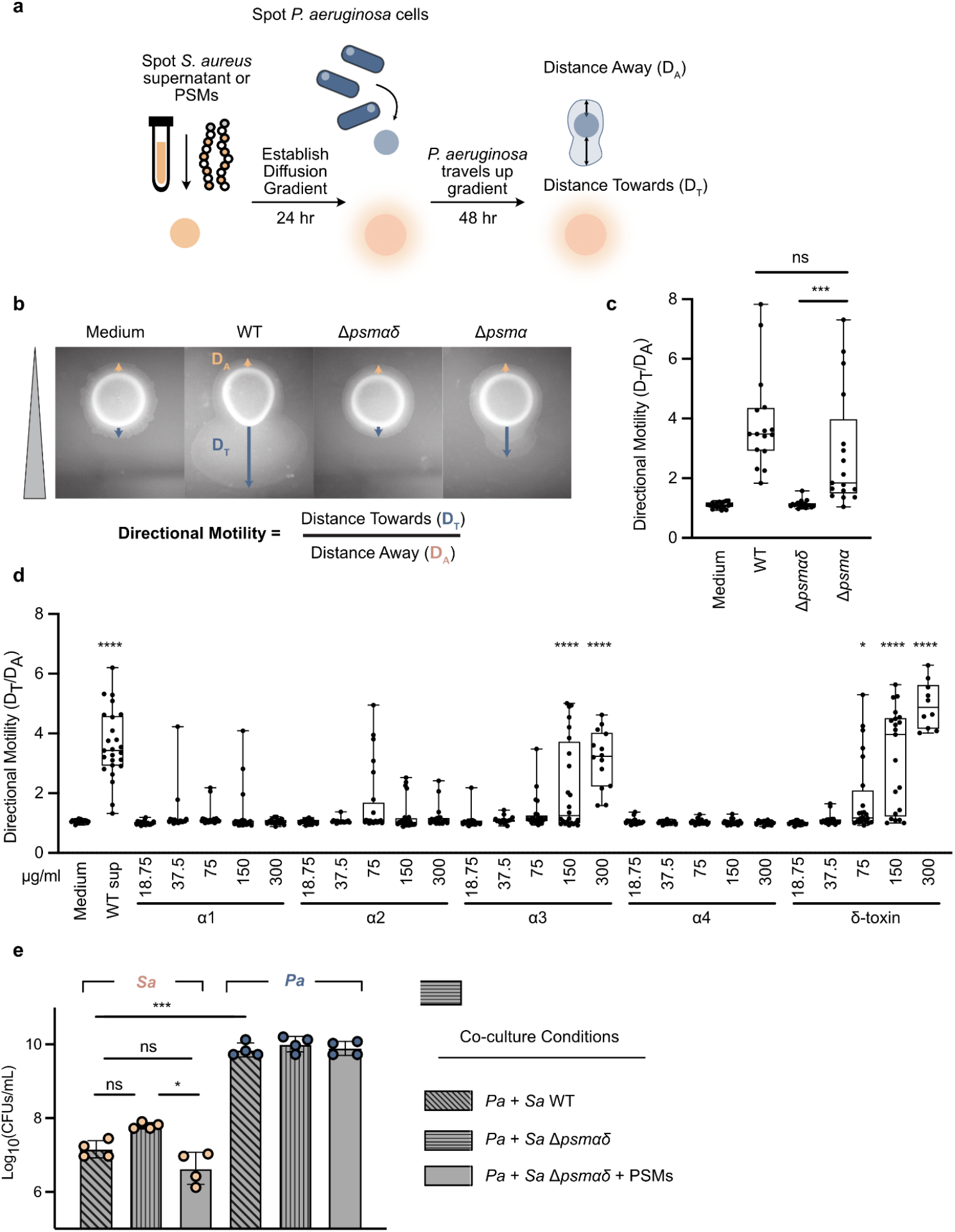
PSMα peptides are necessary and sufficient for *P. aeruginosa* attraction towards *S. aureus*. **a**. Schematic of macroscopic TFP-mediated chemotaxis assays to monitor directional *P. aeruginosa* motility up a pre-established gradient of cell-free *S. aureus* supernatant. Directional motility was calculated as ratio of the motility distance towards (D_T_) over distance away (D_A_) from *S. aureus* supernatant spots. **b**. Representative images of *P. aeruginosa* WT in the presence of a gradient of *S. aureus* growth medium or supernatant derived from the indicated strains (Δ*psmα1–4* and Δ*psmα1–4 δATG-ATT* and Δ*psmα1–4 Δβ1–2 δATG-ATT*). Quantification of directional motility towards a gradient of *S. aureus* supernatant (**c**) or synthetic PSM peptides (**d**) with the median, interquartile range, maximum and minimum indicated for three independent experiments performed in triplicate. Statistical significance was determined by one-way ANOVA followed by Dunnett’s multiple comparisons test. ns, not significant; *, *P* ≤ 0.05; ***, *P* ≤ 0.001; ****, *P* ≤ 0.0001. **e**. CFU enumeration for indicated *P. aeruginosa* and *S. aureus* strains in coculture. Statistical significance was determined by one-way ANOVA followed by Tukey’s multiple comparisons test. ns, not significant; *, *P* ≤ 0.05; ***, *P* ≤ 0.001.

It has been widely acknowledged that *P. aeruginosa* outcompetes *S. aureus in vitro*^13–16^, though the exact mechanisms of cellular death are poorly elucidated. Curiously, when PSM-deficient *S. aureus* were cocultured with *P. aeruginosa*, a moderate increase in *S. aureus* survival was observed (**Fig. 1e**). Addition of PSM peptides to coculture with the Δ*psm* mutant restored *S. aureus* survival to the reduced level seen with WT strains, raising the possibility that there exist unknown PSM-dependent killing mechanisms between *P. aeruginosa* and *S. aureus*. These factors further led us to investigate the roles of PSMs in mediating *P. aeruginosa* responses to *S. aureus*, and the cellular events occurring after *P. aeruginosa* cells are recruited to the site of *S. aureus*.

### *P. aeruginosa* undergoes immediate, systematic proteome remodeling in response to PSM peptide pulse-in and coculture with *S. aureus*

To gain insight into the effects PSMs have on *P. aeruginosa* cellular functions, we took advantage of the precise temporal resolution afforded by <u>B</u>io<u>O</u>rthogonal <u>N</u>on-<u>C</u>anonical <u>A</u>mino acid <u>T</u>agging (BONCAT)^17^ to monitor *P. aeruginosa* immediate protein synthesis in response either to direct addition of PSMs or to coculture with *S. aureus* cells (**Fig. 2a**). *P. aeruginosa* cells constitutively expressing an engineered mutant methionyl-tRNA synthetase (NLL-MetRS) allow for selective metabolic labeling of newly synthesized proteins by the azide-bearing methionine (Met) analog: azidonorleucine (Anl) (**Supplementary Fig. 2**). Downstream chemical enrichment^18^ of labeled proteins enables targeted analysis of nascent *P. aeruginosa* protein synthesis during the Anl labeling period.

**Figure 2.**
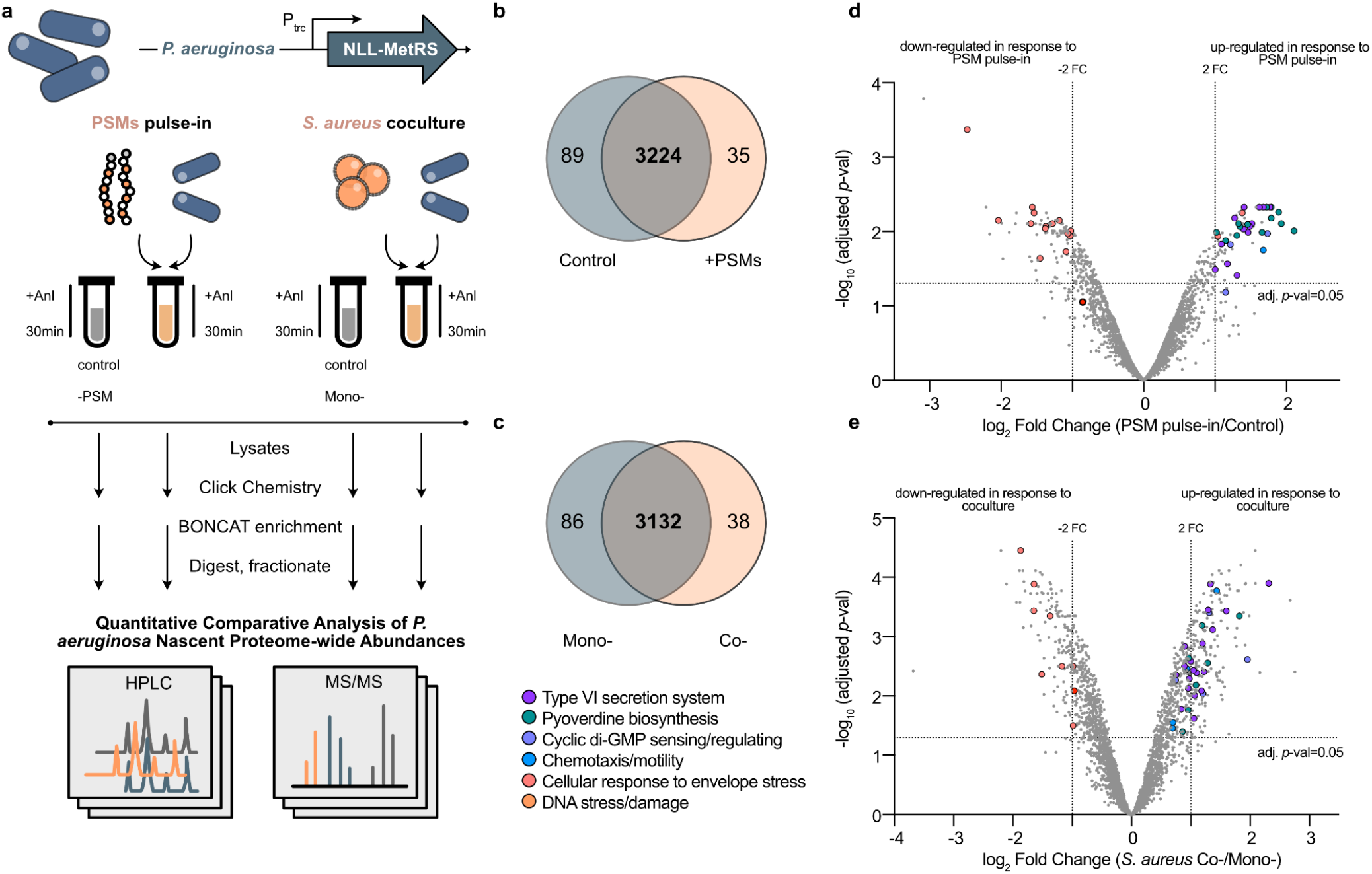
Time-resolved proteome mapping reveals *P. aeruginosa* immediate global responses to PSM peptides pulse-in and coculture with *S. aureus* cells. *P. aeruginosa* cells were engineered to express a mutant tRNA synthetase that allows for metabolic labeling of newly synthesized proteins by a non-canonical amino acid azidonorLeucine (Anl). **a**. Schematic depiction of BONCAT experimental workflow. *P. aeruginosa* protein synthesis immediately following treatment of PSMs pulse-in or coculture with *S. aureus* cells is labeled with Anl for 30 minutes, selectively enriched, and analyzed to compare global nascent proteome abundances with that of untreated control. **b**. Venn diagram showing total proteins quantified for differential expression (overlap) and proteins uniquely identified in +/-PSM pulse-in (**b**) or +/-*S. aureus* coculture (**c**). **d**. Volcano plots summarizing the global proteomic comparisons for +/-PSMs pulse-in conditions and *S. aureus* coculture vs. monoculture conditions (**e**). Protein expression fold-changes between sample groups were calculated via label-free quantification. “Hits” that showed statistically significant changes (Benjamini–Hochberg false-discovery rate adjusted, *P* < 0.05) in abundances in response to PSM pulse-in and *S. aureus* coculture include proteins involved in: type VI secretion system, pyoverdine biosynthesis, c-di-GMP regulation, chemotaxis, motility and cellular responses to envelope stress. n=3 biological replicates for PSM pulse-in proteomics analysis and n=4 for *S. aureus* coculture proteomics analysis.

We identified 3348 and 3365 total proteins newly synthesized by *P. aeruginosa* during the 30-min labeling period immediately following PSM pulse-in and coculture with *S. aureus*, respectively (**Fig. 2b, c**), and quantified differentially expressed proteins in each condition. We found 60 *P. aeruginosa* proteins with statistically significant and greater than 2-fold increase and 98 with greater than 2-fold decrease in abundances in response to PSMs pulse-in. For coculture with *S. aureus* compared to monoculture, 178 proteins with significant increase (>2-fold) in abundances and 124 with significant decrease (>2-fold) in abundances (**Supplementary Table 2**). Candidates were then grouped by their annotated functional categories, which include the following: T6SS, pyoverdine biosynthesis, cyclic di-GMP sensing/regulating enzymes, chemotaxis/motility, cellular response to envelope stress, and DNA damage/stress response (**Fig. 2d, e**). Strikingly, PSMs alone are sufficient to promote the synthesis of proteins in each category.

### *P. aeruginosa* activates T6SS in response to PSMs and *S. aureus* cells

Notably, proteins involved in T6SS are over-represented among the total significantly up-regulated hits in *P. aeruginosa* global proteomic response to PSM pulse-in and *S. aureus* coculture (**Fig. 3, Supplementary Fig. 3**). *P. aeruginosa* T6SS is a speargun-shaped secretory apparatus that loads and injects toxic cargo into prey cells. We detected significantly increased synthesis of various components of the T6SS structural architecture, including core, accessory, bacteriophage-like subunits, and membrane-associated components^19,20^ (**Fig. 3a, c**), suggesting the T6SS apparatus is being systematically assembled during the 30-min labeling period following introduction of PSMs or *S. aureus*. In particular, the expression levels of two proteins—the hemolysin coregulated protein (Hcp, T6SS “sheath”) and the valine-glycine repeat protein G (VgrG, T6SS “tip”)—are often used to determine whether T6SS is functional^5,6,20,21^. Their relative fold-changes are the highest among other T6SS proteins that showed significantly changed abundances in response to PSM pulse-in or coculture with *S. aureus* (**Fig. 3a**). Additionally, proteins encoded by all three known *P. aeruginosa* T6SS loci, denoted HSI-I (PA00-), HSI-II (PA16-) and HSI-III (PA23-)^5,19,20^ were increased (**Fig. 3a, c**), further supporting that *P. aeruginosa* systematically up-regulates T6SS after encountering *S. aureus* via sensing of *Staphylococcal* secreted PSMs.

**Figure 3.**
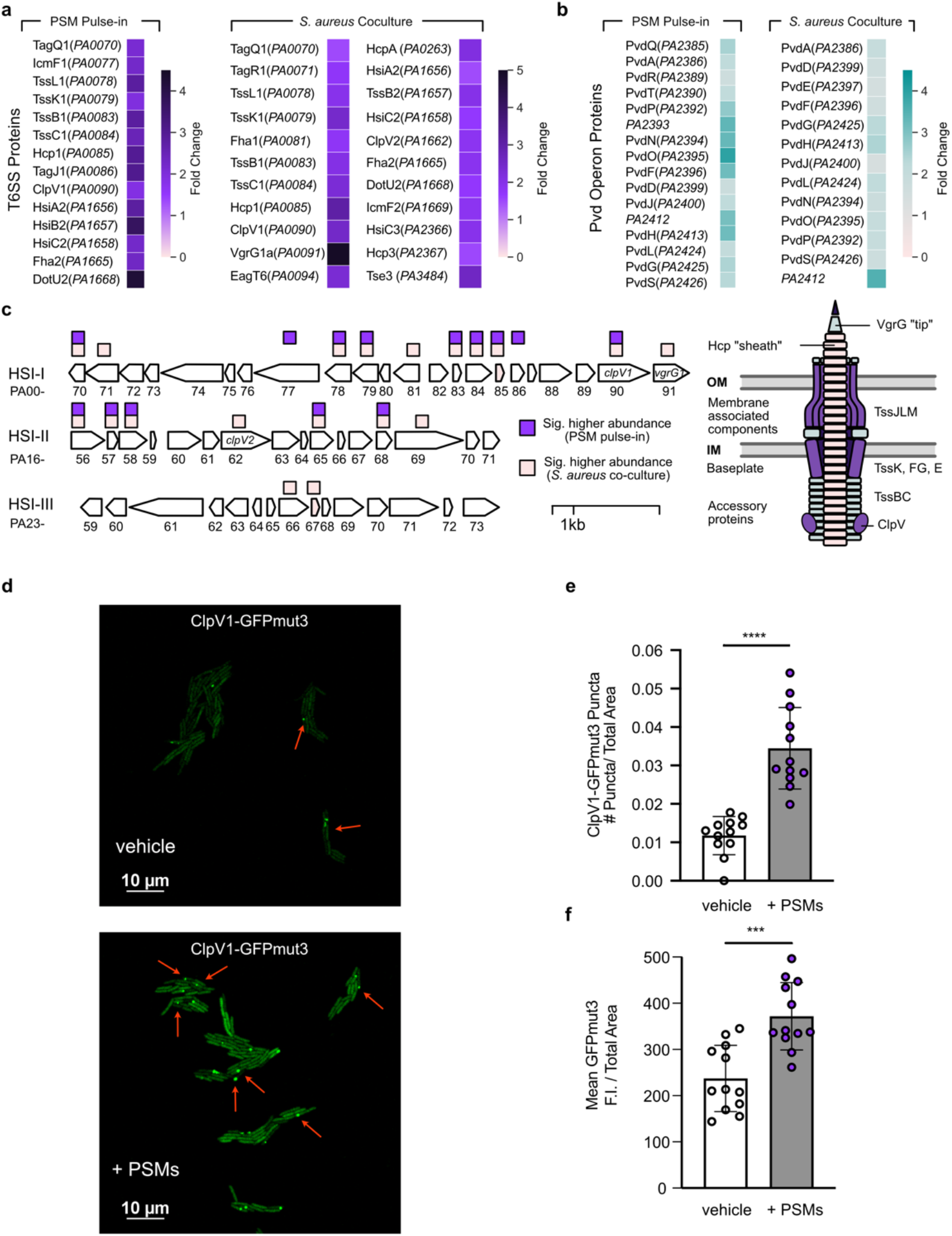
PSMs are interspecies signals that trigger *P. aeruginosa* T6SS antagonism and “competitive stress response”. *P. aeruginosa* induction of T6SS (**a**) and Pvd proteins (**b**) in response to PSMs pulse-in and coculture with *S. aureus* cells. T6SS and pyoverdine biosynthesis proteins with significantly up-regulated fold-changes in response to PSMs pulse-in or *S. aureus* coculture are summarized in associated heatmaps. **c**. Schematic depiction of the *P. aeruginosa* T6SS genetic loci and the 3 HSI-T6SS clusters in *P. aeruginosa* (HSI-I, HSI-II and HSI-III, left) and the structural architecture of the apparatus (right). Squares represent significantly up-regulated in response to PSM pulse-in (purple) or *S. aureus* coculture (peach). **d**. Representative microscopy images of *P. aeruginosa* ClpV1-GFPmut3 fluorescence with vehicle control (water, top) and with synthetic PSM treatment (8 μg/mL, bottom). Examples of ClpV1 fluorescent puncta formation are highlighted with arrows (red). Quantification of number of ClpV1-GFPmut3 fluorescent puncta per cellular total area is shown in (**e**) and mean GFPmut3 fluorescence intensity (F.I.) per cellular total area in (**f**). Data represent a total of three biological replicates with four technical replicates (FOVs) per condition, per biological replicate, analyzed. Statistical significance was determined by unpaired *t-* test: ***, *P* = 0.0001; ****, *P* ≤ 0.0001.

We next ranked the nascent *P. aeruginosa* proteome by individual protein raw abundances quantified by label-free quantification (LFQ) via mass spectrometry (**Supplementary Fig. 4**) to examine cellular allocation of protein synthesis resources following PSM pulse-in and *S. aureus* coculture challenge. Remarkably, most T6SS proteins appeared in the top quartile with significantly elevated average abundances in PSM-treated and *S. aureus* coculture samples compared to untreated/monoculture controls, further indicating T6SS antagonism is prioritized by *P. aeruginosa* in responding to interspecies stress.

Although T6SS apparatus assembly does not necessarily indicate firing of T6SS effectors, significantly higher abundances of the AAA+ ATPases ClpV (**Fig. 3a, c**) suggest increased sheath contraction and propulsion of effectors^22-24^. To examine *P. aeruginosa* deployment of T6SS, single-cell microscopy using a fluorescent reporter of ClpV1 activity was employed (ClpV1-GFPmut3^9^) and confirmed that PSMs are sufficient to induce *P. aeruginosa* deployment of T6SS (**Fig. 3d**). PSM-treated cells exhibited both significantly increased GFPmut3 puncta formation (**Fig. 3e**) as well as overall fluorescence intensity (**Fig. 3f**) per cellular total area, further supporting that PSMs induce interspecies antagonistic T6SS attacks by *P. aeruginosa*.

### *Staphylococcal* secreted PSM peptides increase siderophore biosynthesis

We also observed that the T6SS induction in *P. aeruginosa* is accompanied by systematic upregulation of the pyoverdine biosynthesis cluster (**Supplementary Fig. 3**), which produces a siderophore that binds to extracellular Fe^3+^ with high affinity^25,26^. Iron starvation is a major stress response pathway evolutionarily conserved in bacteria. Proteins encoded by five pyoverdine operons (**Fig. 3b**) entirely covering the complex cellular biosynthesis machinery for pyoverdine siderophore were found to be significantly up-regulated in response to PSM pulse-in or *S. aureus* coculture—including the extracytoplasmic function iron starvation σ factor PvdS, which positively regulates pyoverdine biosynthesis and secretion^27^, and PvdR, which controls transport of pyoverdine out of the cell^28^, indicating that siderophores are being increasingly synthesized and dispatched out of the cell during the 30-min labeling period. Thus, we simultaneously monitored pyoverdine production and induction of gene expression using a fluorescent reporter P’*pvdG*-*mScarlet*^29^ (**Supplementary Fig. 5**). Consistent with the proteomic results, we observed significantly increased pyoverdine production by *P. aeruginosa* following PSMs treatment, as well as significant induction of *pvdG* promoter activity.

Pathogens face intense competition for iron with host and other microbial species due to the essentiality of iron as a nutrition source, and siderophore production is often reported to be involved in exploitive interspecies competition^13,30,31^. Interestingly, a recent study reported upregulation of siderophore biosynthesis in *P. aeruginosa* when treated with *Staphylococcal* culture supernatant^29^, though the molecular signals responsible for the observed upregulation remained elusive. Here, we show that *Staphylococcal* secreted PSM peptides alone could trigger increased pyoverdine biosynthesis and export, further suggesting that PSMs play important roles in mediating interspecies competition between *P. aeruginosa* and *S. aureus*.

### PSMs may activate competition sensing via induction of transient membrane stress

We next probed the molecular mechanism of PSM-induced T6SS activation. Previous literature suggests *P. aeruginosa* T6SS could be induced via kin cell lysis^9^ and/or envelope stress^32^. In particular, the pore-forming antibiotic polymyxin B induces T6SS in *P. aeruginosa* via endogenous membrane stress^32^. *S. aureus* secreted PSMs are virulence factors with hemolytic activity toward mammalian cells^11^. While PSMs generally exhibit low activity towards bacterial membranes^33^, we asked whether PSMs could permeabilize the *P. aeruginosa* membrane, cause kin cell lysis, and/or cause envelope stress in *P. aeruginosa*. Live imaging of *P. aeruginosa* with propidium iodide +/-PSMs did not show evidence of kin lysis, inner membrane permeability (**Fig. 4a**), or altered *P. aeruginosa* growth rate (**Supplementary Fig. 6a**). In comparison, polymyxin B significantly inhibited *P. aeruginosa* growth (**Supplemental Fig. 6a**) and induced a moderate uptake of propidium iodide (**Fig. 4a**). Further analysis of outer membrane permeability by uptake of 1-N-phenylnaphthylamine (NPN) also did not reveal significant permeability with PSM treatment, while polymyxin significantly induced outer membrane permeability (**Fig. 4b**).

**Figure 4.**
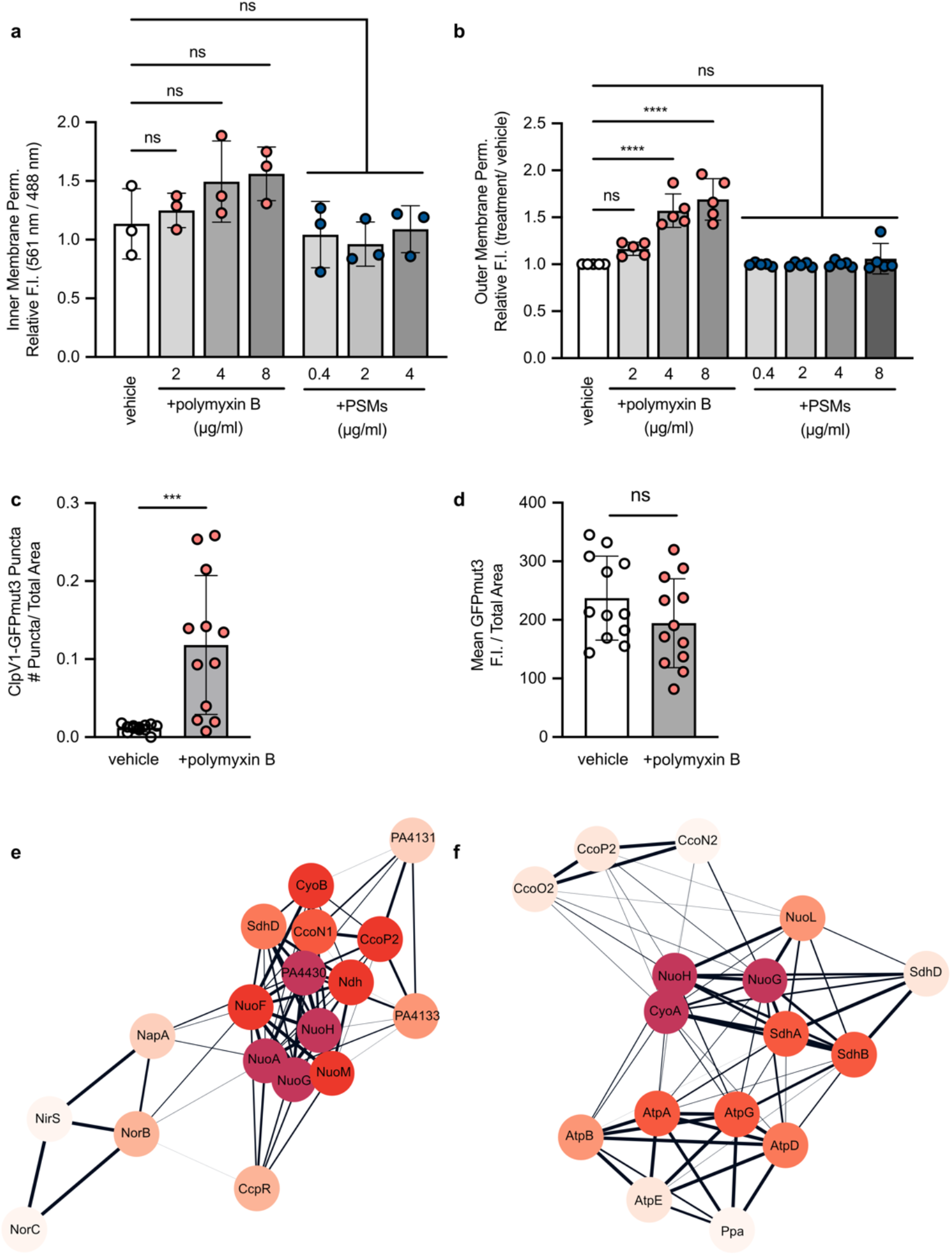
PSMs may activate competition sensing via induction of transient membrane stress. **a**. Inner membrane permeability was determined by calculating the ratio of propidium iodide fluorescence to GFP fluorescence of PA14 P*tac-gfp* exposed to the indicated total concentrations of polymyxin B or synthetic PSMɑ1 and PSMɑ3. **b**. Outer membrane permeability was determined by measuring NPN fluorescence of *P. aeruginosa* cells incubated with polymyxin B or PSMs. Data shown represent the mean and standard deviation of at least three independent experiments. For **a**-**b**, statistical significance was determined by one-way ANOVA followed by Dunnett’s multiple comparisons test. ***, *P* ≤ 0.001; ****, *P* ≤ 0.0001. **c**. Quantification of number of ClpV1-GFPmut3 fluorescent puncta per cellular total area following treatment with polymyxin B. **d**. Mean GFPmut3 fluorescence intensity (F.I.) per cellular total area following treatment with polymyxin B. For **c-d**, Data represents a total of three biological replicates with four technical replicates (FOVs) per condition, per biological replicate, analyzed. Statistical significance was determined by unpaired *t-*test: ns, not significant; ***, *P* ≤ 0.001. See representative microscopy images in **Supplementary Fig. 6b**. STRING protein interaction network for ETC proteins with significantly decreased abundances in PSM pulse-in (**e**) and in response to *S. aureus* coculture (**f**).

Given these differences in membrane activity between polymyxin B and PSMs, we revisited T6SS activation by polymyxin B with ClpV1 fluorescent reporter^9^ under the current study conditions for comparison. Polymyxin B-treated cells displayed distinct ClpV1 puncta induction (**Fig. 4c**) but yielded low mean fluorescence intensity per cell (**Fig. 4d**), suggesting potential molecular differences between mechanisms of T6SS induction by polymyxin B versus by PSMs treatment. Polymyxin B can be inserted into the membrane, causing cell lysis by creating pores in the envelope^34^. In contrast, PSMs are cationic, amphipathic small helical peptides with membrane perturbing and cell surface-adhering properties^35^.

While we were unable to detect significant membrane damaging activity by PSMs, we hypothesize that non-lethal membrane perturbations may explain PSM-induced T6SS activation in *P. aeruginosa*. Several factors contributed to this hypothesis: first, global differential proteomic profiling revealed significant and systematically decreased production of electron transport chain (ETC) enzymes in response to PSMs pulse-in and coculture with *S. aureus* (**Fig. 4e, f, Supplementary Fig. 3**), a characterized cellular response to envelope stress evolutionarily conserved in *E. coli* and other Gram-negative bacteria^36–39^. In addition, while we did not detect increased protein synthesis of classic regulators of membrane stress, such as σ^E^ and CpxAR^39^, we observed significant up-regulation of a subset of proteins involved in membrane stress responses, most notably protein encoded by *PA3731*, a close homologue of the phage shock protein PspA in *E. coli* and member of a family of proteins characterized to play crucial roles in the cellular response to and protection against envelope stress in *E. coli* and other Gram-negative species^40,41^. Therefore, we hypothesize that PSMs provoke *P. aeruginosa* T6SS firing via induction of cell envelope stress via short-term perturbations.

### Significantly increased *P. aeruginosa* T6SS activity in coculture with *S. aureus* on CF patient-derived bronchial epithelial cells

Previous studies reported that Hcp1 is detected at high levels in chronic CF sputum^5^, and HSI-II and III T6SS are required for and induced upon *P. aeruginosa* infection of epithelial cells^7,42^, suggesting that differential regulation of any of the three T6SS loci in polymicrobial infections may have implications for the host. Prompted by the fact that all three HSI-T6SS loci in *P. aeruginosa* have previously characterized roles in CF pathogenicity, we further investigated *P. aeruginosa* and *S. aureus* interactions in a host-derived environment to explore interspecies virulence factor crosstalk in a clinically relevant context.

For this purpose, we obtained the health care-associated methicillin-resistant *Staphylococcus aureus* (HA-MRSA) strain USA100, a highly antibiotic resistant clinical isolate and a leading cause of invasive infections by MRSA^43,44^, and *P. aeruginosa* strain PAO1, a laboratory derivative more closely related than PA14 to most clinical isolates of CF^45^. We performed RNA-sequencing to examine *P. aeruginosa* transcriptomic changes that contribute to interspecies interactions in a coinfection model with *S. aureus* using polarized, fully differentiated CF bronchial epithelial cells (CFBE41o-, **Fig. 5a**). This model closely mimics the CF host environment by recapitulating approximately 84% of *P. aeruginosa* gene expression in human expectorated CF sputum, outperforming both laboratory media and the acute mouse pneumonia model of infection^46^.

**Figure 5.**
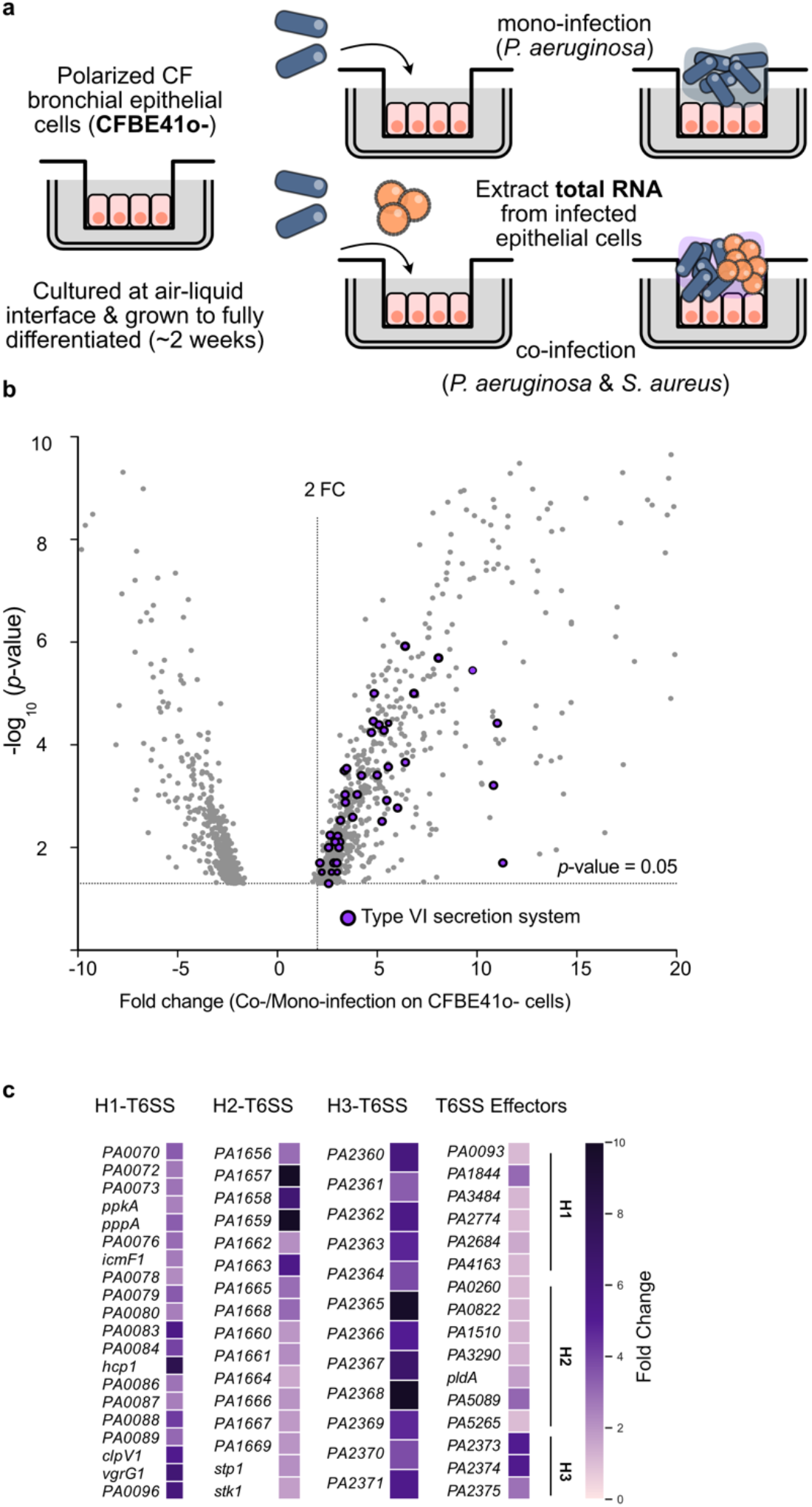
*P. aeruginosa* T6SS activity is significantly increased during coculture with *S. aureus* on CF patient-derived bronchial epithelial cells. **a**. Schematic of dual-species RNA-seq approach. CF bronchial epithelial cells (CFBE41o-) were seeded at air-liquid interface and allowed to fully differentiate. Polarized cells were infected apically with *P. aeruginosa* alone or cocultured with *S. aureus* for 6 hours before total RNA was collected. **b**. Volcano plot summarizing differentially expressed *P. aeruginosa* genes in coculture with *S. aureus* as compared to *P. aeruginosa* mono-infection. **c**. Heatmaps summarizing significant differential fold changes of T6SS genes grouped by the three HSI-T6SS clusters. Data represent the mean fold change from two independent, biological replicates.

We identified 1,325 differentially expressed genes during coculture with *S. aureus* (fold change >2 or <-2, P<0.05). Of these, we detected increased transcription of T6SS genes from all three HSI-T6SS clusters (**Fig. 5b, c**). Transcription of *hcp, vgrG*, and sheath genes was significantly increased, suggesting that the T6SS apparatus is functional. In addition, we observed significantly increased transcription of several effector genes including *tse1* (*PA1844*), a peptidoglycan amidase^47^, *pldB* (*PA5089*), a phospholipase^48^, and *tseF* (*PA2374*)^49^, a known facilitator of iron uptake in *P. aeruginosa*^48^.

### *P. aeruginosa* T6SS mediates the killing of *S. aureus*

A wealth of existing literature shows that T6SS-delivered effectors target and kill Gram-negative bacteria^9,10,20,47^. However, until a recent study that demonstrated T6SS secreted effectors by *Acinetobacter baumannii* could kill Gram-positive bacteria^50^, it had previously been assumed that Gram-positive species are not susceptible to T6SS-mediated killing^51^.

To examine if T6SS activity provides *P. aeruginosa* a competitive advantage in coculture with *S. aureus* in association with airway epithelial cells, we constructed clean deletions of each HSI T6SS sheath gene, ∆ *tssB1* (H1), ∆ *hsiB2* (H2), and ∆ *hsiB3* (H3), and surprisingly found that each mutant exhibited decreased competitive index compared to WT in coculture with *S. aureus* in the airway cell model (**Fig. 6a, b**). We next focused on a HSI-III T6SS-encoded effector TseF for several reasons. First, *tseF* (*PA2374*) was the most significantly up-regulated effector in coinfection with *S. aureus* on host cells (**Fig. 5c**). Secondly, TseF facilitates *P. aeruginosa* iron uptake^49^, a functional role likely to affect polymicrobial competition. Further, TseF was characterized to be coregulated with the *Pseudomonas* quinolone signal^49^, a quorum-sensing system in *P. aeruginosa* with known roles in competition against *S. aureus*^15^. Interestingly, deletion of *tseF* alone significantly reduced *P. aeruginosa* competitive fitness against *S. aureus* and rescued *S. aureus* survival in coculture (**Fig. 6a, b**).

**Figure 6.**
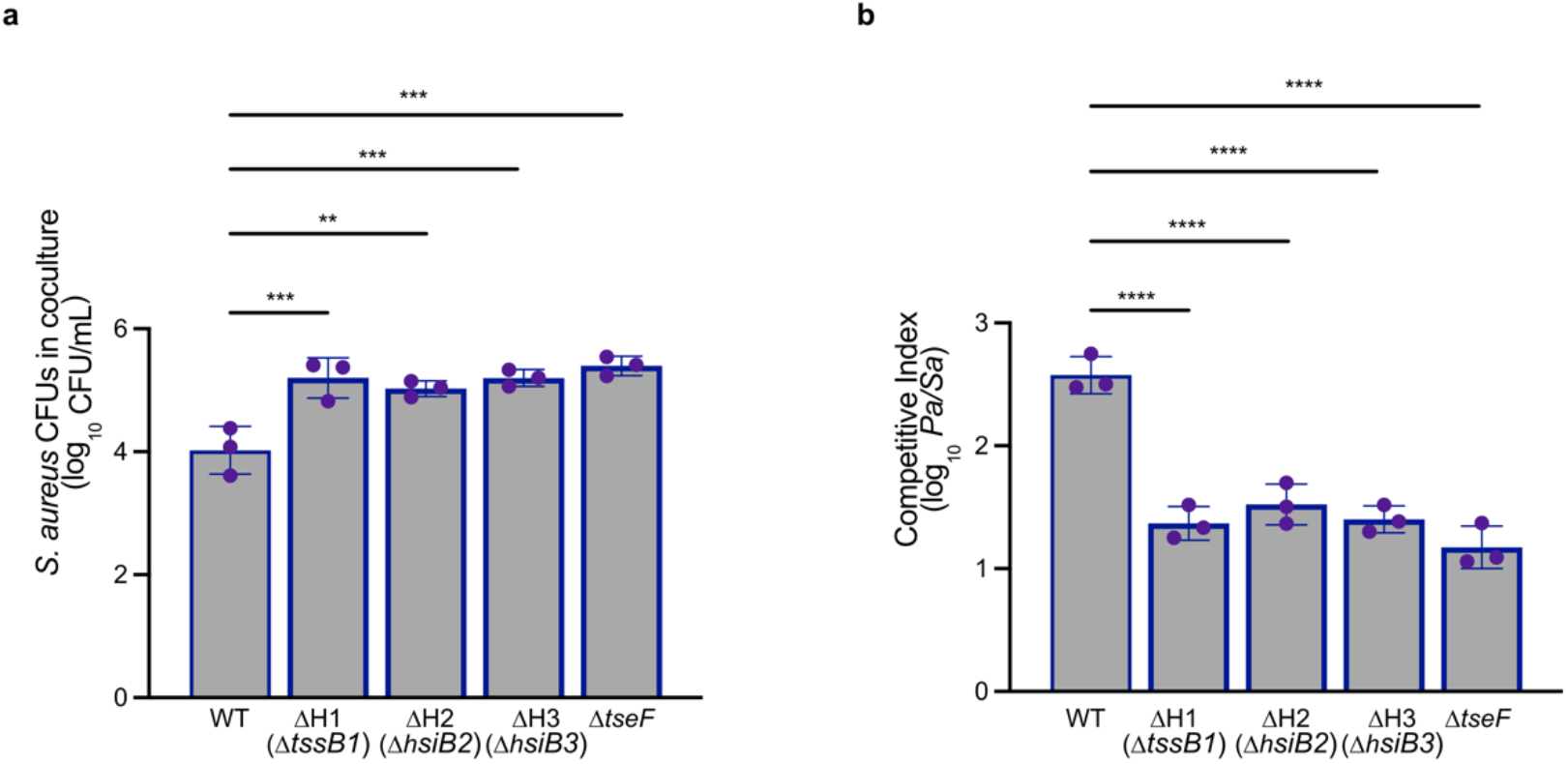
*P. aeruginosa* T6SS contributes to competition with *S. aureus*. Coculture CFU enumeration of *S. aureus* in coculture with *P. aeruginosa* on CFBE41o-cells. **a**. *S. aureus* survival **b**. *P. aeruginosa* competitive indexes (log_10_ (*Pa* CFU / *Sa* CFU) for indicated mutant strains. Data represent three independent biological replicates. Statistical significance was determined by one-way ANOVA followed by Dunnett’s multiple comparisons test. **, *P* ≤ 0.01; ***, *P* ≤ 0.001; ****; *P* ≤ 0.0001.

## DISCUSSION

The “competition-sensing” hypothesis states that bacteria adapt evolutionarily conserved stress response pathways to directly detect and respond to ecological competition^52^. The results presented here provide empirical evidence for this hypothesis, which predicts increased bacterial toxin production in response to stress caused by competitors. We propose a model in accordance, whereby *P. aeruginosa* senses transient cellular stress caused by secreted competitor signals and swiftly responds by moving towards the signals and activating antagonistic responses (**Fig. 7**). Activation of membrane stress and iron starvation responses observed in *P. aeruginosa* further supports that “competition sensing” is manifested in several stress response pathways.

**Figure 7.**
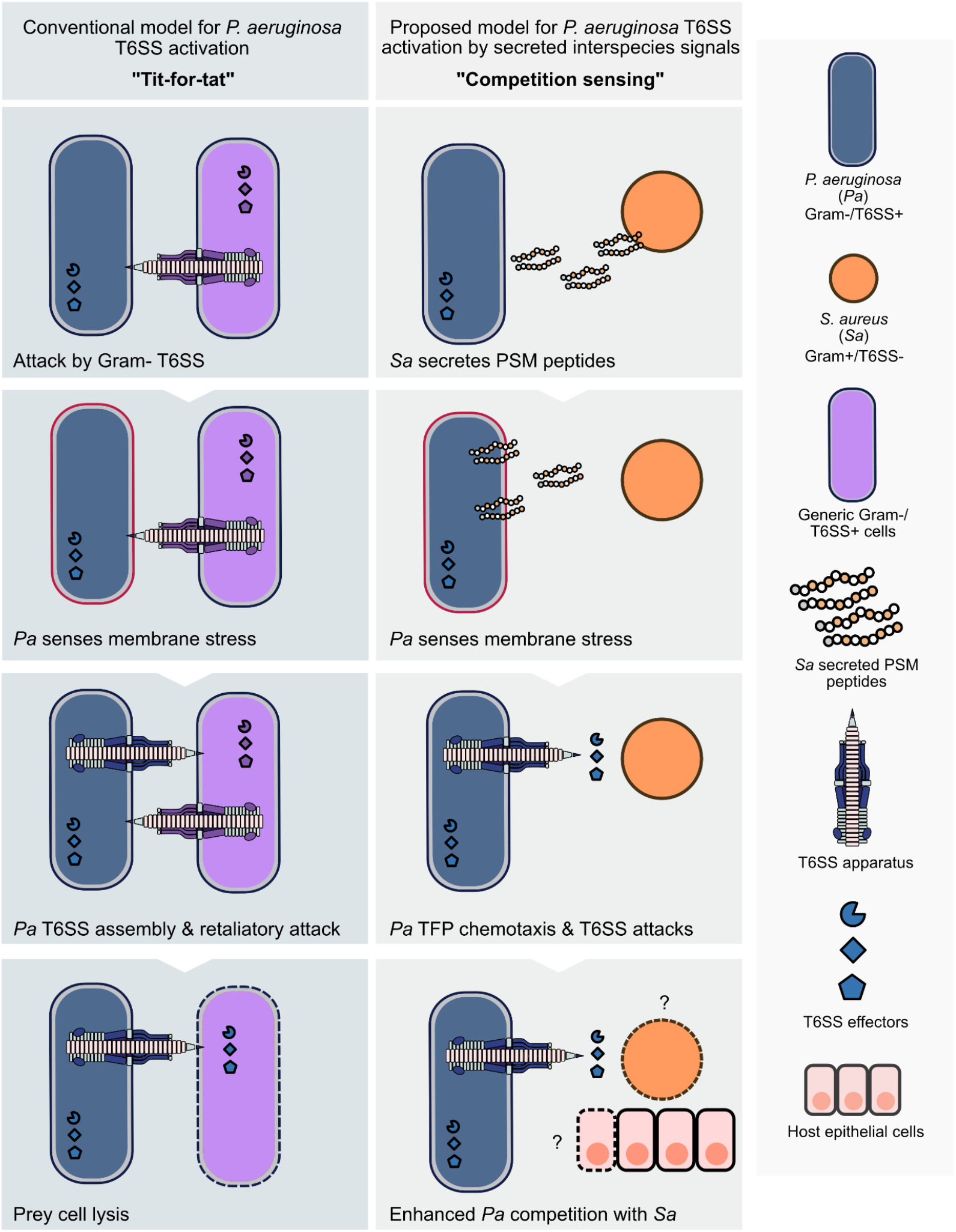
Proposed model for *P. aeruginosa* T6SS activation by *S. aureus* secreted interspecies signals. *P. aeruginosa* has been characterized to activate its T6SS following exogenous attacks in a “tit-for-tat” pattern (left panel), assembling the T6SS apparatus and preferentially firing at other Gram-negative, T6SS-positive species. T6SS “dueling” is known to result in the lysis and death of the competitor (left). In “competition sensing” (right panel), *P. aeruginosa* detects *Staphylococcal* secreted PSM peptides (potentially through membrane stress), subsequently travels towards *S. aureus* with increased directional motility and simultaneously activates T6SS firing. *P. aeruginosa* T6SS activation is sustained during coinfection with *S. aureus* and results in enhanced competitive fitness against *S. aureus*. Effectors may be delivered by T6SS apparatus or secreted extracellularly and have bactericidal or static action towards *S. aureus*. Increased *P. aeruginosa* T6SS activity, particularly of the HSI-II and III, during coinfection may have detrimental consequences on host epithelial cells and contribute to worsened CF patient clinical outcome.

*P. aeruginosa* is attracted to diverse bacterial species and moves towards the site of competition^5^; thus, a close analogy can be drawn between *P. aeruginosa* and the notorious predacious bacterium, *Myxococcus xanthus*, characterized to coordinate group responses to invade and lyse prey^53^. *P. aeruginosa* displays incipient multicellularity via complex collective behaviors, including ones of a predatory nature as described here. We propose that upon sensing interspecies signals, *P. aeruginosa* cells move to “trap” a *S. aureus* colony, further enabling contact-dependent invasion and/or local concentration of secreted antimicrobials.

One potential mechanistic model of competition sensing is that *P. aeruginosa* closely monitors cell envelope integrity to detect environmental and/or interspecies insults. While PSMs do not affect the *P. aeruginosa* membrane sufficiently to allow permeabilization, even transient envelope stress may induce T6SS assembly and firing. Interestingly, it has been recently reported that *P. aeruginosa* chemotaxis towards, instead of away from, antibiotics and releases bacteriocins before dying^54^. While PSMs did not reduce *P. aeruginosa* viability, we found induction of two pyocins in *P. aeruginosa* in response to both PSMs pulse-in and co-culture with *S. aureus*, potentially supporting a similar “suicidal chemotaxis” model. PSMs alone are sufficient to trigger TFP-mediated motility, synthesis and transport of siderophores, activation of T6SS antagonism and envelope stress responses, suggesting that PSMs are important interspecies signals that help *P. aeruginosa* sense and respond to imminent danger/competition. Interestingly, T6SS, pyoverdine production, chemotaxis and cellular response to envelope stress in *P. aeruginosa* are all known to be regulated by cyclic di-GMP^55–59^. We propose herein that secondary messengers signaling networks mediate “competition sensing” and global bacterial responses to interspecies insults. In support of this hypothesis, we observed up-regulation of multiple c-di-GMP metabolizing enzymes, suggesting several c-di-GMP mediated signaling networks are activated and are involved in *P. aeruginosa* response to PSMs and *S. aureus* (**Supplementary Fig. 7**).

Further, proteomic analysis detected significantly increased abundance of PA1611, a known inhibitor of RetS and activator of the <u>g</u>lobal <u>a</u>ctivation of antibiotic and <u>c</u>yanide synthesis/regulator of <u>s</u>econdary <u>m</u>etabolism (Gac/Rsm) pathway^59^ both in response to PSM treatment and *S. aureus* coculture. Gac/Rsm post-transcriptionally regulates all three T6SS loci in *P. aeruginosa*^61^ and mediates *<u>P</u>. <u>a</u>eruginosa* response to <u>a</u>ntagonism (PARA)^10^. Also consistent with previous reports that *P. aeruginosa* T6SS and T3SS are inversely regulated via RetS^56^, we detected systematic repression of T3SS and simultaneously increased T6SS activity during coinfection with *S. aureus* (**Supplementary Fig. 8**). Intriguingly, the Gac/Rsm pathway and c-di-GMP signaling networks both regulate T6SS and iron uptake^20,58,59^. Future work will be dedicated to studying overlap in signal transduction pathways and potential coordination of interspecies phenotypes reported in this study, including *P. aeruginosa* TFP-mediated directional motility, downstream antagonistic attacks and exploitive iron scavenging.

Interestingly, we observed inverse regulation of siderophore biosynthesis in coculture with *S. aureus* using global proteomics analysis performed *in vitro* versus transcriptomic analysis performed in a host environment. *P. aeruginosa* down-regulates pyoverdine biosynthesis during coinfection with *S. aureus* on CF-derived epithelial cells (**Supplementary Fig. 8**). We attribute this to the differences in temporal resolution of the experiments—while chemo-selective proteomic analysis captured immediate “competition sensing” responses, global RNA-sequencing investigated long-term coinfection phenotypes. These results highlight *P. aeruginosa* versatile genetic plasticity in regulating iron scavenging behaviors during short-term versus long-term competition and underline the importance of studying and comparing polymicrobial interactions both *in vitro* and *in vivo*.

Numerous studies have reported that *P. aeruginosa* produces diverse secondary metabolites known to be toxic to *S. aureus*^14,15^, but insufficient to account for total *S. aureus* cellular death in coculture^16^. Nonetheless, when embarking on this study, we presumed that *P. aeruginosa* T6SS would neither be activated by, nor effective in competition with *S. aureus*. Several reasons contributed to this initial assumption^51^. First, Gram-positive bacteria lack a conjugative pilus, and therefore cannot provoke *P. aeruginosa* reciprocal firing. Further, the Gram-positive cell wall constitutes a thicker peptidoglycan (PG) layer in comparison to that of Gram-negative species, which was thought to prohibit penetration by the T6SS apparatus and effective delivery of toxic effectors. The discovery here that *P. aeruginosa* T6SS is both induced by and mediates the killing of a Gram-positive pathogen, challenges our prior assumptions, and expands the role of T6SS during infection, opening a wealth of new opportunities to study, inhibit, or co-opt interspecies competition.

How does *P. aeruginosa* T6SS kill *S. aureus*? Intriguingly, proteomic, transcriptional, and mutational analyses suggest that all three HSI loci have a role in facilitating the killing of *S. aureus*. While we focused on the HSI-III T6SS effector TseF for further study due to its significantly increased transcript level revealed by RNA-sequencing analysis, future work will be dedicated to defining the scope and specificity of functionality for *P. aeruginosa* antagonism against *S. aureus* mediated by T6SS effectors. Moreover, the global proteomics study was only performed on the *P. aeruginosa* intracellular lysate fraction, which did not include most secreted protein effectors found in the extracellular fraction; thus, it is possible *S. aureus* induces the secretion of T6SS effectors not identified here. A recent study demonstrated that Tse4, a T6SS muramidase effector of *A. baumannii*, exhibits promiscuous PG-degrading activity and kills Gram-positive species, including *S. aureus*^50^. While previous literature indicated T6SS-exported muramidases generally cannot effectively lyse Gram-positive cells^62^, the possibility remains that certain PG-targeting T6SS effectors can impact cellular functions of Gram-positive bacteria, not limited to causing cellular death or lysis. Beyond cell wall degrading toxins, developing evidence that suggests the T6SS apparatus can inject and deliver effectors into the Gram-positive cell wall^50^ points to the emerging possibility that diverse T6SS effectors could have bacteriostatic and bactericidal potential towards both Gram-negative and Gram-positive bacteria. For instance, studies analyzing differential regulation for *S. aureus* in coculture with *P. aeruginosa* consistently reported up-regulation of SOS response and oxidative stress response pathways^15,40^, but it remained unclear how *P. aeruginosa* triggers these responses in *S. aureus*. It is therefore curious to speculate that these effects could be due to previously unknown attacks by *P. aeruginosa* antibacterial T6SS nuclease toxins^63^ and NAD(P)+ glycohydrolases effectors^64^.

Cumulatively, our findings provide a new model of T6SS-mediated interspecies interactions for Gram-negative and Gram-positive species. Our results revealed complex polymicrobial virulence factors crosstalk and highlight the importance of leveraging a comprehensive molecular understanding of polymicrobial competition while studying the host-pathogen interface. Considering both *Staphylococcal* PSMs and *P. aeruginosa* T6SS have well-characterized functions in modulating host immune responses, their interactivity uncovered by our study could have detrimental implications on the host.

## Acknowledgements

This work was supported by the Jacobs Institute for Molecular Engineering for Medicine and the Center for Environmental Microbial Interactions at Caltech, and by the Institute for Collaborative Biotechnologies through cooperative agreement W911NF-19-2-0026 from the U.S. Army Research Office, the Cystic Fibrosis Foundation (LIMOLI19R3 to DHL and BOMBER18G0 to JMB), and the National Institutes of Health (1R35GM142760-01 to DHL and 1R01HL142587 to JMB). We thank Drs. Megan Bergkessel (University of Dundee), Melanie Spero (University of Oregon), Alex Horswill (University of Colorado Denver), Mike Schurr (University of Colorado Denver), and Li Wu (University of Iowa) for helpful discussions and valuable insight. We also thank members of the Limoli and Tirrell Labs for careful editing of the manuscript and helpful discussions. We thank Dr. Jeff Jones (Caltech) for an in-house pipeline for proteomics data processing, Dr. J. Muse Davis for the use of the stereoscope, and Drs. Joseph Mougous and Anupama Khare for the generous gifts of the ClpV1-GFPmut3 and P’*pvdA-mScarlet* reporters, respectively.

## Competing interests

The authors declare no competing interests.

## Notes

### Competing Interest Statement

The authors have declared no competing interest.

