## Supplemental text and figures for "*Staphylococcal* secreted cytotoxins are competition sensing signals for *Pseudomonas aeruginosa*"

### Materials and Methods

#### Bacterial strains, plasmids and culture conditions

*P. aeruginosa* and *Escherichia coli* were routinely cultured in lysogeny broth (LB; 1% tryptone, 0.5% yeast extract, 1% sodium chloride) and *S. aureus* in tryptic soy broth (TSB, Becton Dickenson), unless otherwise indicated, with aeration at 37ºC. Antibiotics were added, when appropriate: carbenicillin at 250 µg/ml for *P. aeruginosa* and gentamycin (Gm) at 30 µg/ml for *P. aeruginosa* and 15 µg/ml for *Escherichia coli.* All strains used in this study can be found in **Table 1**.

#### Generating *P. aeruginosa* mutants

Markerless deletion mutations were constructed in PAO1 to inactivate each of the *P. aeruginosa* T6SS loci. Deletions were introduced using two-step allelic exchange, as previously described for *P. aeruginosa*[1]. Briefly, 500 bp upstream and downstream regions of the *tssB* gene from each T6SS loci (H1-H3-T6SS) were amplified to construct a contiguous mutant allele by SOE-PCR. The deletion allele was inserted into the donor vector pENTRPEX18Gm by Gateway cloning, transformed into *E. coli* S17.1, and introduced to *P. aeruginosa* through conjugation using a mating filter apparatus. Deletion mutants were first confirmed by PCR and followed by whole genome sequencing and *breseq* analysis to ensure no secondary mutations arose. All oligonucleotides used to generate *P. aeruginosa* mutants can be found in **Table 1**.

#### Macroscopic coculture twitching chemotaxis assay

Motility experiments were performed as previously described[2–4]. Buffered agar plates (10 mM Tris, pH 7.6; 8 mM MgSO4; 1 mM NaPO4, pH 7.6; and 1.5% agar) were poured and allowed to solidify for 1 hour prior to incubation for 16 hours at 37°C and 22% humidity. After solidifying, 4 µL of either growth medium (TSB) or cell-free supernatant derived from an overnight culture of USA300 *S. aureus* at $OD_{600}$ 5.0 and filter sterilized with a 0.22 µm filter were spotted on the surface of the plate and allowed to diffuse for 24 hours at 37°C and 22% humidity to establish a gradient. *P. aeruginosa* PA14 cultures were incubated overnight in TSB with aeration at 37°C, subcultured 1:100 in TSB the following morning, then standardized to $OD_{600}$ 12.0 in 100 µL of 1 mM MOPS buffer supplemented with 8 mM $MgSO_4$ prior to inoculating 1 µL on the surface of the plate at five mm from the center of the gradient. Plates were incubated in a single layer, agar-side down, for 24 hours at 37°C with 22% humidity, followed by an additional 16 hours at room temperature prior to imaging the motility response of *P. aeruginosa.* Images were captured using a Zeiss stereoscope with Zeiss Axiocam 506 camera and directional motility ratios were calculated in Fiji before graphing and performing statistical analysis in GraphPad Prism.

**BONCAT labeling of *P. aeruginosa***

For PSM pulse-in and *S. aureus* coculture proteomics experiments, *P. aeruginosa* PA14 NLL-MetRS cells were grown to $OD_{600}$~2 in defined M14 medium (M9 salts supplemented with 10 mM glucose, 10 mg/ml casamino acids, 1 mM $MgSO_4$, 2 µg/ml thiamine (vitamin B1), 2 µg/ml niacin (vitamin B3), 2 µg/ml calcium pantothenate (vitamin B5), 0.1 µg/ml biotin (vitamin B9)). For PSM pulse-in, PSMα1 and PSMα3 (Genscript, 0.2 µg/ml each) were added to *P. aeruginosa* cells; for cocultures, *P. aeruginosa* (PA14 NLL-MetRS) and *S. aureus* (USA300_FPR3757) cells were mixed at 1:1 ratio. Labeling was initiated by the addition of 1 mM Anl (Iris-Biotech) and cell lysates were harvested immediately after 30 min. Cell pellets were centrifuged at 4°C and washed with ice-cold LCMS-grade water and frozen at -80°C for downstream lysis and chemical enrichment. All samples were lysed by resuspension in 1% SDS in PBS, boiled at 75°C and sonicated with a microtip probe for 15 s at 30% amplification (Qsonica).

**Sample preparation for mass spectrometry**

For chemical enrichment of BONCAT-labeled proteins from lysates, 2.5 mg of lysates were used for each sample. Lysates were first reduced by dithiothreitol (DTT), alkylated with chloroacetamide and reacted with aza-dibenzocyclooctyne (DBCO) agarose beads (Click Chemistry Tools) via copper-free click chemistry. The reaction was incubated in dark at room temperature overnight, followed by rigorous resin washing with i) 40 ml 0.8% (wt/vol) SDS/PBS; ii) 40 ml 8 M urea in tris hydrochloride (pH = 8.0) and iii) 40 ml 20% (vol/vol) acetonitrile/$H_2O$ to remove nonspecific protein binding. Peptides were detached from DBCO agarose with on-bead digestion with 0.1 $\mu$g trypsin and 0.05 $\mu$g endoproteinase LysC at 37°C overnight. Supernatant was then collected for further detergent removal (HiPPR column, Thermo Fisher Scientific), and peptides were desalted with $C_{18}$ StageTips.

**LC-MS analyses of desalted samples**

Peptides were subjected to LC-MS/MS analysis on an EASY-nLC 1200 (Thermo Fisher, San Jose, CA) coupled to a Q Exactive HF Orbitrap mass spectrometer (Thermo Fisher, Bremen, Germany) equipped with a Nanospray Flex ion source. Samples (2 µL out of 10 µL total) were directly loaded onto an Aurora 25 cm x 75 µm ID, 1.6 µm C18 column (Ion Opticks, Victoria, Australia) heated to 50°C. The peptides were separated with a 60 min gradient at a flow rate of 350 nL/min as follows: 2–6% Solvent B (3.5 min), 6-25% B (42 min), 25-40% B (14.5 min), 40-98% B (1 min), and held at 98% B (14 min). Solvent A consisted of 97.8% H2O, 2% ACN, and 0.2% formic acid and solvent B consisted of 19.8% H2O, 80 % ACN, and 0.2% formic acid. The Q Exactive HF was operated in data dependent mode with full scan resolution set to 60,000 at m/z 200 in profile mode, full scan target set to $3 \times 10^6$, and a maximum injection time of 15 ms. Full scan mass range was set to 375−1500 m/z. Data dependent MS2 scans were collected for charge state 2-5 precursors in centroid mode using a loop count of 12, AGC target of 1 × $10^5$, an intensity threshold of 1 × $10^5$, and a maximum injection time of 45 ms. Isolation width was set at 1.2 m/z, scan range was 200-2000 m/z, and a fixed first mass of 100 was used. Normalized collision energy was set at 28. Peptide match was set to off, and isotope exclusion was on. Dynamic exclusion was set to exclude after 1 time for 45 sec. All mass spectrometry raw data used for this study have been deposited to the California Institute of Technology Research Data Repository for public access under DOI 10.22002/gbjeb-6kg53.

**Proteomics data processing and analysis**

Proteomics data analysis was performed in Proteome Discoverer 2.5 (Thermo Scientific) using the SequestHT search algorithm and a nonredundant Uniprot *P. aeruginosa* FASTA file plus common contaminants (CRAPome). Sequest search parameters were as follows: fully tryptic peptides with 6-144 residues and no more than 2 missed cleavages, precursor mass tolerance of 35 ppm and fragment mass tolerance of 0.05 Da, and a maximum of 3 equal modifications. Dynamic modifications were as follows: Cysteine carbamidomethylation, Methionine oxidation, Asparagine and Glutamine deamidation, protein N-terminal acetylation, protein N-terminal Met-loss, and protein N-terminal Met-loss plus acetylation. Percolator FDRs were set at 0.01 (strict) and 0.05 (relaxed). The consensus level peptide and PSM FDR filters were also set at 0.01 (strict) and 0.05 (relaxed). Strict parsimony principle was set to true. Protein quantification was also performed in Proteome Discoverer by summed abundances of the precursor intensities of all high confidence unique and razor peptides.

Raw protein quantification data exported from ProteomeDiscoverer 2.5 was imported into R and analyzed using the romics analysis package (https://jeffsocal.github.io/romics). Once imported the data is filtered for common protein contaminants and normalized between runs using Breiman and Cutler's Random Forests for Classification and Regression (https://cran.r-project.org/web/packages/randomForest) method selected based on best performance in lowering sample replicate variation while maintaining quantitative dynamic range for label free analysis. Protein expression differences between samples were evaluated in the R romics package using the limma algorithm for differential expression (https://bioinf.wehi.edu.au/limma/). Differential expression volcano plots were generated using Prism (GraphPad). For PSM pulse-in proteomics analysis, principal component analysis revealed an outlier for PSM-treated samples (data not shown). Differential expression analysis performed before ($log_2$ fold change and raw P-values) and after ($log_2$ fold change and Benjamini Hochberg FDR adjusted P-values) removal of outlier yielded consistent observations, and the final volcano plots shown represent data analysis post removal of the outlier. Protein set enrichment analysis for significantly up-regulated and down-regulated *P. aeruginosa* proteins was performed using STRING v11 database (Search Tool for Retrieval of Interacting Genes/Proteins) by uploading protein accession codes (PAO1 UniProt) to the server (https://string-db.org/), and results visualized using python libraries matplotlib and networx.

**Fluorescence microscopy**

To visualize the ClpV1 foci, *P. aeruginosa* PAO1 ClpV1-GFPmut3 was grown in minimal medium (M8 salts supplemented with 0.2% glucose and 1.2% tryptone, M8T) with aeration at 37°C. Cells were sub-cultured in fresh M8T, grown to mid-log phase ($OD_{600}$ of ~0.3 - 0.6), standardized to $OD_{600}$ of 0.3, pelleted, and resuspended in M8T (vehicle) or 8 µg/ml PSMs (4 µg/ml α1 and 4 µg/ml α3). PSMs were sonicated for 5 min prior to use. 0.5 µL of cells were inoculated onto a 4-chamber glass-bottom 35 mm dish (Cellvis, Cat. no. D35C4-30-1.5-N) before placing an agarose pad on top. Agarose pads were made by pipetting 550 µL of M8T with 2% molten agarose (Lonza, Cat. no. 50081) into each quadrant of a 4-chamber glass-bottom dish and drying uncovered for ~1 hr 15 min at room temperature, followed by ~1 hr covered with a lid at room temperature, then ~1 hr 15 min at 37°C before transferring the pads onto the inoculated glass-bottom dish. For polymyxin B, since significant cellular toxicity was observed at all concentrations examined, which also increased ClpV1-GFP puncta formation when added at $T_0$, a final concentration of 4 µg/ml of polymyxin B was added on top of the agarose pad ~2 hrs after the cells were inoculated, and then allowed to diffuse for ~40 min. The imaging was performed with an inverted Nikon Ti2 A1R Galvanometer Scanning Confocal Microscope, using a 100x Plan Apo oil objective (1.45 NA). Images were acquired between 2 – 3 hrs after the cells were inoculated onto the dish. A 488 nm laser was used to excite GFPmut3. Images were saved and analyzed using the Nikon NIS-Elements AR software and the representative images are shown in the Maximum Intensity projection. The ClpV1-GFPmut3 puncta were manually quantified and divided by the total cellular area per field of view (FOV). The mean fluorescence intensity (F.I.) was quantified by calculating the average of the mean F.I. of the total cellular area per FOV. A total of three biological replicates with four technical replicates (FOVs) per condition, per biological replicate, were analyzed.

For determination of inner membrane permeability to propidium iodide, *P. aeruginosa* cells harboring P*tac-gfp* on a plasmid were grown in M8T with 250 µg/ml carbenicillin to mid-log phase and prepared as described above except 0.5 µM propidium iodide (PI) was added to the agarose pads and polymyxin was added directly to the cells at $T_0$. Phase contrast and epifluorescent images were acquired with an Andor Sona camera. Fluorescent images were acquired every 20 minutes with TxRed images taken at 18 ms exposure and 20% Sola fluorescent light and GFP imaged at 2 ms exposure and 20% Sola fluorescent light. The rate of PI uptake was determined by dividing the total fluorescence intensity of PI by the total fluorescence intensity of GFP in each FOV. A total of three biological replicates with two technical replicates (FOVs) per condition, per biological replicate, were analyzed.

**Outer membrane permeability**

*P. aeruginosa* cells were grown in M14 at 37°C with aeration to early stationary phase and standardized to $OD_{600}$ = 1.0. Cells were then washed in fresh M14 and incubated with 20 µM 1-N- Phenylnaphthylamine (NPN) and polymyxin B or synthetic PSMα1 and PSMα3 for 15 minutes. 100 µL from each condition were added to a black, clear-bottom 96 well plate and NPN fluorescence read (excitation: 350 nm, emission: 420 nm) on a Tecan Infinite M200 plate reader. Relative NPN fluorescence was calculated by dividing fluorescence of treated cells by fluorescence of untreated cells.

***P. aeruginosa* growth rate**

*P. aeruginosa* cells were grown in M14 to mid-log phase, washed in fresh M14, standardized to $OD_{600}$ = 0.05, and added to a clear-bottom 96-well plate with medium alone, polymyxin B, or synthetic PSMα1 and PSMα3. The plate was covered with a gas permeable membrane and $OD_{600}$ measurements were taken at 10 min intervals on a Tecan infinite M200 plate reader. The plate was incubated at 37°C and was shaken for 1 s before each measurement.

**Mammalian cell culture**

The human CF bronchial epithelial cell line CFBE41o- (obtained from J. P. Clancy, Cincinnati Children's Hospital) originated from a person with CF that was homozygous for the ΔF508 mutation in CFTR. CFBE41o- cells were routinely cultured in minimal essential media (MEM) supplemented with 10% fetal bovine serum (FBS), 2 mM L-glutamine, 5 U/ml penicillin-5 mg/ml streptomycin, and 0.5 mg/ml plasmocin at 37°C with 5% CO2. CFBE41o- cells were seeded at near confluency on Transwell filters (Costar). After confluency, CFBE41o- cells were differentiated at an air-liquid interface (ALI) for 1-2 weeks before use in experiments. The identity and purity of the CFBE41o- cells were verified by short tandem repeat profiling (University of Arizona Genetics Core). Cells were tested quarterly for mycoplasma using a Southern Biotech mycoplasma detection kit.

**Coinfection of CFBE cells**

ALI-differentiated CFBE41o- cells were moved to antibiotic-free media by washing with MEM (Gibco) supplemented with 2 mM L-glutamine and replacing basolateral media with MEM supplemented with 10% FBS and 2 mM L-glutamine. *P. aeruginosa* PAO1 and *S. aureus* USA100 were pre-washed in MEM supplemented with 2 mM L-glutamine and inoculated at a 1:1 ratio at an MOI of approximately 250 onto the apical side of polarized CFBE41o- cells. After 2 hours of attachment, non-attached bacteria were removed, and apical media was adjusted to 0.4% L-arginine for a total of 6 hours.

**RNA sequencing**

RNA was isolated from *P. aeruginosa* PAO1 and *S. aureus* USA100 mono- or 1:1 coculture following 6-hour infection on polarized CFBE41o- cells. RNA was collected by phenol:chloroform extraction using RNA-Bee (AMS Biotechnology) and zirconia/silica beads in a BeadBeater (BioSpec Products) from two independent, biological replicates. RNA was precipitated with isopropanol and linear acrylamide, and RNA pellets were washed by ethanol precipitation. RNA was treated with Turbo DNase (Ambion) and purified by RNA Clean and Concentrator (Zymo Research). DNA removal was confirmed by 260/280 and 260/230 ratios and by PCR for the *P. aeruginosa rplU* gene. RNA integrity was determined by agarose gel electrophoresis and visualization of 5S, 16S, 18S, 23S, and 28S bands.

RNA-seq library preparation and sequencing were performed by the Health Sciences Sequencing Core at Children's Hospital of Pittsburgh (Pittsburgh, PA). RNA concentration and integrity was confirmed by fluorometric quantification (Qubit) and Tapestation analysis (Agilent). RNA was rRNA-depleted using Ribo-Zero Epidemiology, and sequencing libraries were prepared using Truseq stranded Total RNA Kit (Illumina). Single end sequencing was performed on a NextSeq 500. Approximately 75 million 75 bp reads were obtained for each sample. Reads were processed and mapped to the *P. aeruginosa* PAO1 genome and differential expression analysis was performed in CLC Genomics Workbench. Statistically significant changes were considered as $p \leq 0.05$. The *P. aeruginosa* and *S. aureus* co-infection transcriptome sequencing (RNA-seq) data is part of a larger study on the response of *P. aeruginosa to co-infection*. This data has been deposited in the NCBI Sequence Read Archive (BioProject #PRJNA865724).

**Bacterial competition assay**

For *in vitro* coculture competition, *P. aeruginosa* (PA14) and *S. aureus* (USA300 WT and Δ*psm*α1-4 δATG-ATT) overnight cultures were diluted to $OD_{600}$ = 0.1 in fresh M14 medium, mixed at 1:1 ratio by species and grown for 24 hr. PSMα1 and PSMα3 (Genscript, 5 μg/ml each) were added to the medium at time of inoculation of coculture. Harvested cocultures were serially diluted and plated on *Pseudomonas* isolation agar (PIA) for selective isolation of *P. aeruginosa* and Mannitol Salt Agar for selective isolation of *S. aureus*.

For bacterial competition associated with CFBE41o- cells, *P. aeruginosa* (PAO1) was grown overnight in LB and *S. aureus* (USA100) was grown overnight in TSB at 37˚C. Overnight cultures were washed with phosphate buffered saline (PBS). Strains were OD-normalized and mixed at a 1:1 ratio. As before, differentiated and polarized CFBE41o- cells were infected at an MOI ~250 with the 1:1 bacterial mixture. After 1 hour of attachment to the airway cells, planktonic bacteria were removed, and the competition assay was run for a total of 6 hours. Following competition, 0.1% Triton-X was added to the apical compartment to remove attached cells and enumerate bacteria. For each assay, the initial 1:1 inoculum (input) and 6-hour time point (output) were serially diluted in PBS and spotted on PIA to count *P. aeruginosa* and on tryptic soy agar supplemented with 10 mg/L colistin sulfate (Sigma) and 15 mg/L nalidixic acid (Sigma) to count *S. aureus*. Colony forming units (CFUs) were determined for both species, and competitive index was calculated as: $[(WT_{output}/competitor_{output})/(WT_{input}/competitor_{input})]$.

### Extended Data Figures

**a**

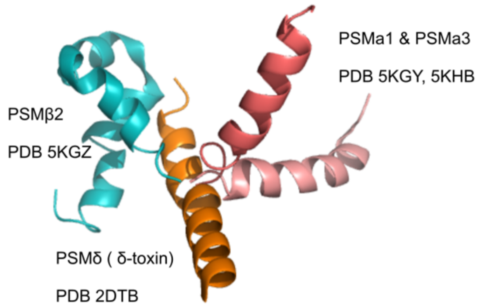


δ-toxin: fMAQDIISTIGDLVKWIIDTVNKFTKK (26)
PSMα1: fMGIIAGIIKVIKSLIEQFTGK (21)
PSMα2: fMGIIAGIIKFIKGLIEKFTGK (21)
PSMα3: fMEFVAKLFKFFKDLLGKFLGNN (22)
PSMα4: fMAIVGTIIKIIKAIIDIFAK (20)
PSMβ1: fMEGLFNAIKD TVTAAINNDG AKLGTSIVSI VENGVGLLGK LFGF (44)
PSMβ2: fMTGLAEAIAN TVQAAQQHDS VKLGTSIVDI VANGVGLLGK LFGF (44)

**b**

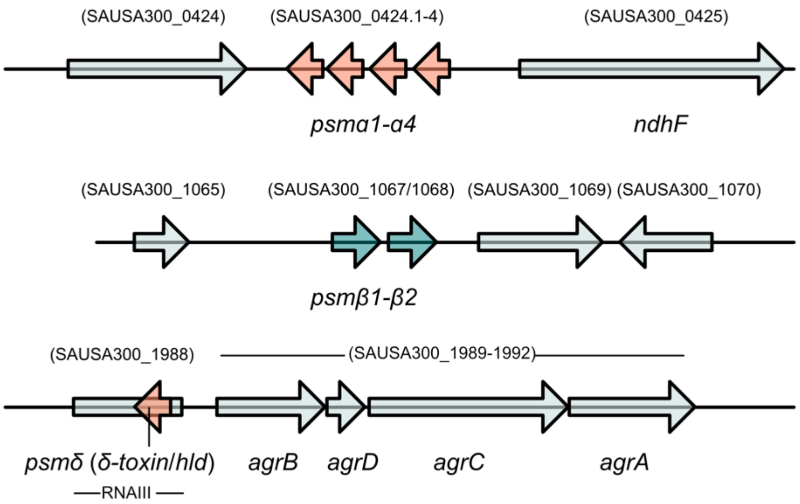


**Supplementary Figure 1. PSM peptide sequences, structure and genome location.**

**a.** Amino acid sequences for all 7 known peptides in the PSM family of *Staphylococcus aureus* with deposited Protein Data Bank (PDB) structures shown. f=N-formylation. **b.** Regulatory operons and gene locations for PSMs in *S. aureus* (annotations based on strain USA300 FPR3757). Adapted from[5]. The *psmα1-4* mutant used previously[11], was generated via deletion of the entire *psmα* operon (*psmα1 - α4*). To generate the double *psmαδ* mutant, since *psmδ* is encoded within the coding region of RNAIII, *psmδ* was inactivated through mutation of the start codon from ATG to ATT[11].

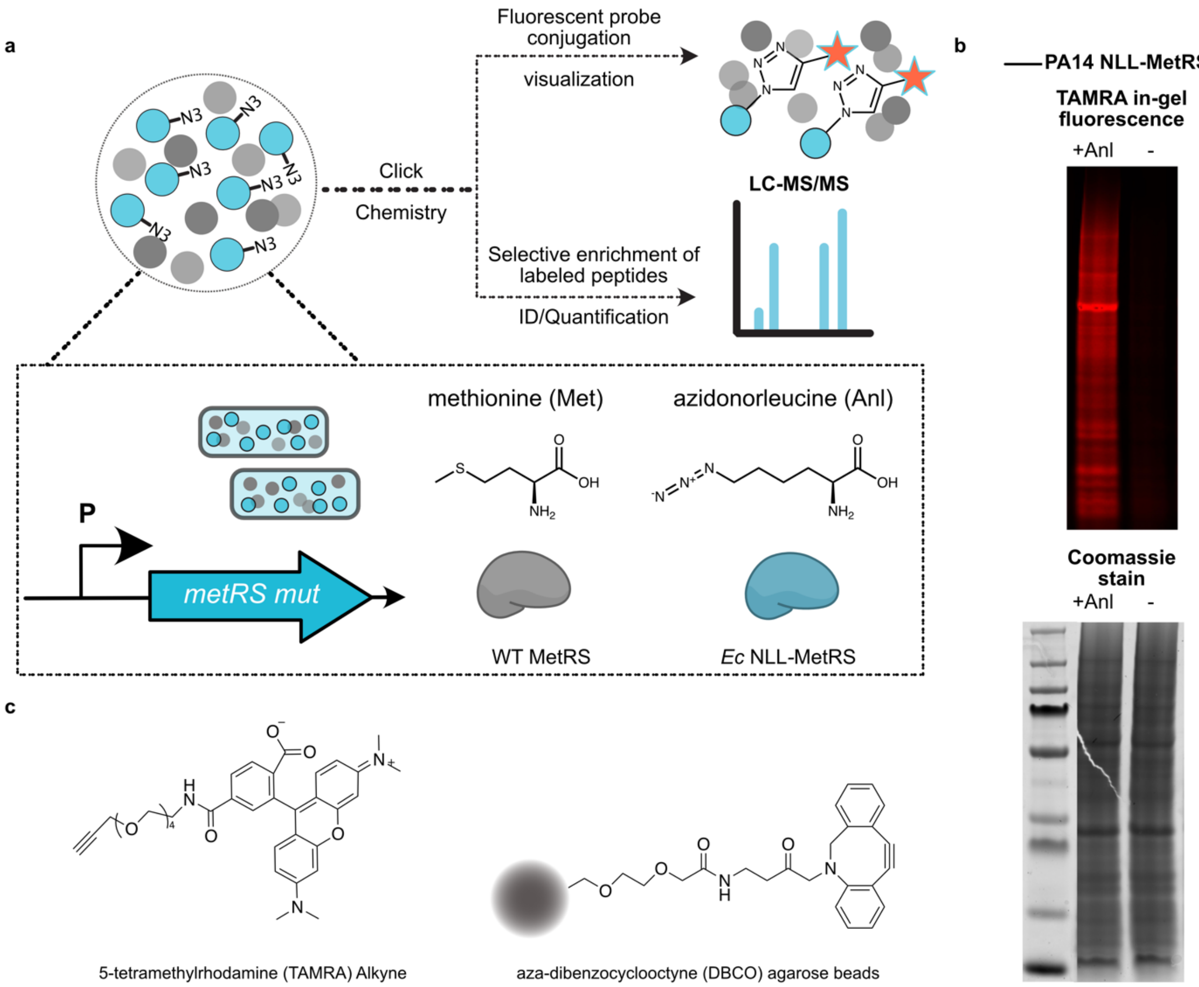


**Supplementary Figure 2. BONCAT labeling and enrichment of nascent *P. aeruginosa* proteome.** **a.** General scheme of a BONCAT experiment. *P. aeruginosa* cells constitutively expressing NLL-MetRS were treated with 1 mM azidonorleucine (Anl) to initiate protein labeling. Newly synthesized proteins (blue circles) were selectively tagged with azide functional groups on Anl residues and chemically distinct from the pre-existing cellular proteome (grey circles). Labeled proteins were enriched via copper-free azide-alkyne click chemistry onto DBCO-alkyne agarose beads. Enriched proteins were digested and analyzed by LC-MS/MS or visualized via in-gel fluorescence following copper-catalyzed azide-alkyne click chemistry onto TAMRA-alkyne fluorophore (**b**). Anl-labeled proteins in each sample were detected and visualized by TAMRA fluorescence (552 nm ex/ 578 nm em) gel via copper-catalyzed click chemistry as previously described[6]. Briefly, lysates were quantified by the Pierce$^{TM}$ BCA Protein Assay kit (Thermo Fisher Scientific), and samples containing 100 $\mu$g protein were incubated with 2.5 $\mu$M alkyne-TAMRA (Click Chemistry Tools), 100 $\mu$M $CuSO_4$, 500 $\mu$M tris(3-hydroxypropyltriazolylmethyl)amine (THPTA ligand; Click Chemistry tools) and 5 mM aminoguanidine hydrochloride. The click reaction was initiated by addition of 5 mM sodium ascorbate and allowed to proceed for 30 min in the dark at room temperature, followed by methanol/chloroform precipitation. Precipitated proteins in each sample were then washed twice with methanol, resuspended in PBS and SDS loading buffer, and visualized via SDS-PAGE electrophoresis (NuPAGE Novex 4-to12% bis-Tris gels; Thermo Fisher Scientific). Imaging was done on a Typhoon gel imager (GE Healthcare) for fluorescence or coomassie (Instant-Blue) signal intensities. **c.** Chemical compounds used for BONCAT in-gel fluorescence and enrichment in this study.

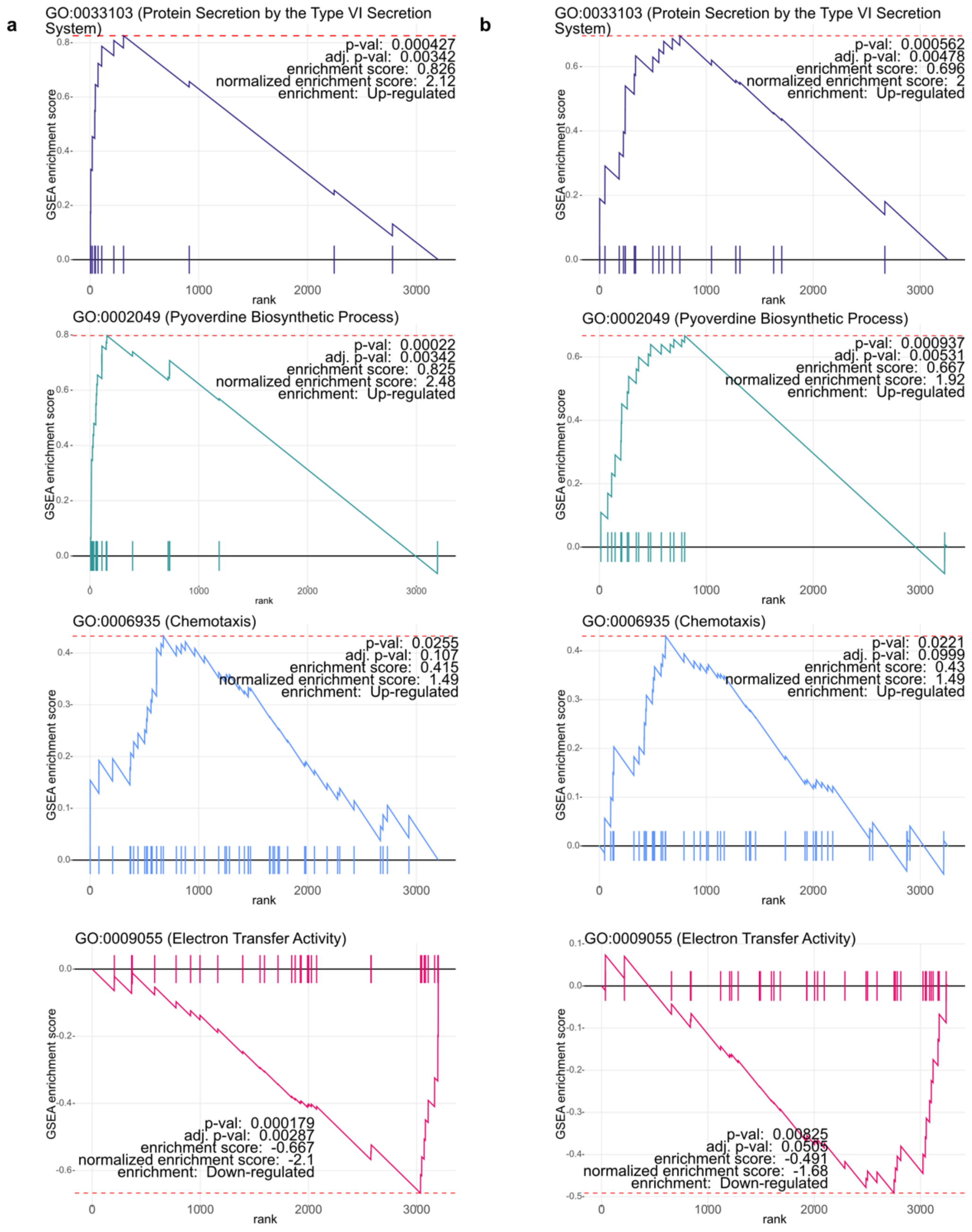


**Supplementary Figure 3. Enrichment analysis of proteins differentially regulated in response to PSM pulse-in and *S. aureus* coculture.**

Gene set enrichment analysis (GSEA)[7] was performed for **PSM pulse-in (a) and *S. aureus* coculture (b)** using the fgsea package for R (https://bioconductor.org/packages/release/bioc/html/fgsea/html). GSEA compares a rank-ordered list of genetic and/or proteomic data to curated annotations. Briefly, proteins with quantified differential expressions were provided as a rank-ordered list by $log_2$-transformed fold changes with annotations pulled from the *Pseudomonas* Genome Project (pseudocap.gaf) and downloaded UniProtKB entry annotations for *P. aeruginosa*. Analysis revealed significantly enriched Gene Ontology (GO) terms for cellular components, molecular functions and biological processes with indicated false discovery rate (FDR) adjusted *P* values. The number of permutations was set to 10,000 for *P* value calculation. For complete lists of GSEA output, see **Supplementary Table 2** and **3**.

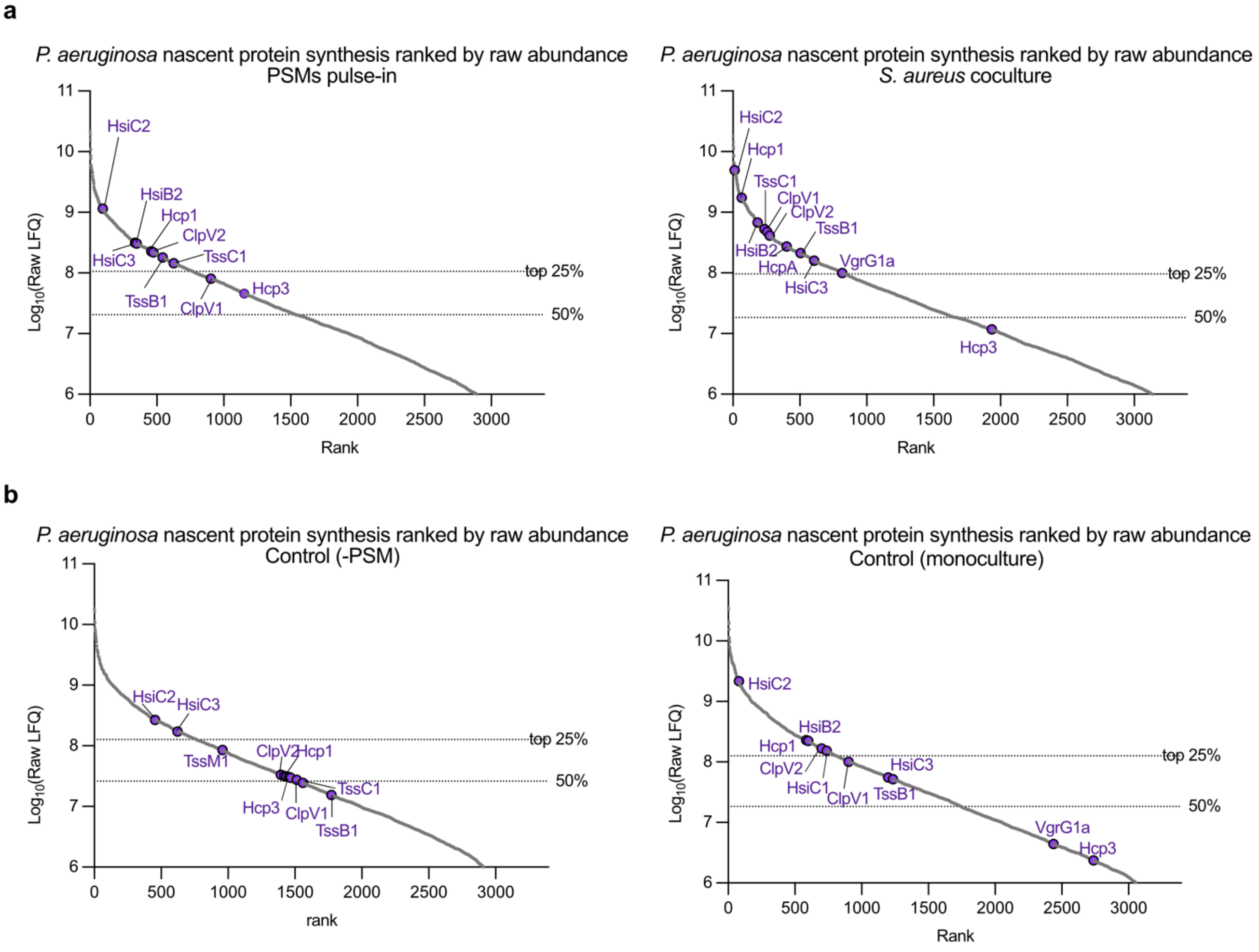


**Supplementary Figure 4. *P. aeruginosa* proteome ranked by average raw abundances (T6SS core components highlighted).**

**a.** *P. aeruginosa* nascent protein synthesis in response to PSM pulse-in and *S. aureus* coculture during Anl labeling period is ranked by individual protein raw abundance as indicated by Label-Free Quantification (LFQ) values. **b.** *P. aeruginosa* nascent protein synthesis in control (-PSM and monoculture) samples.

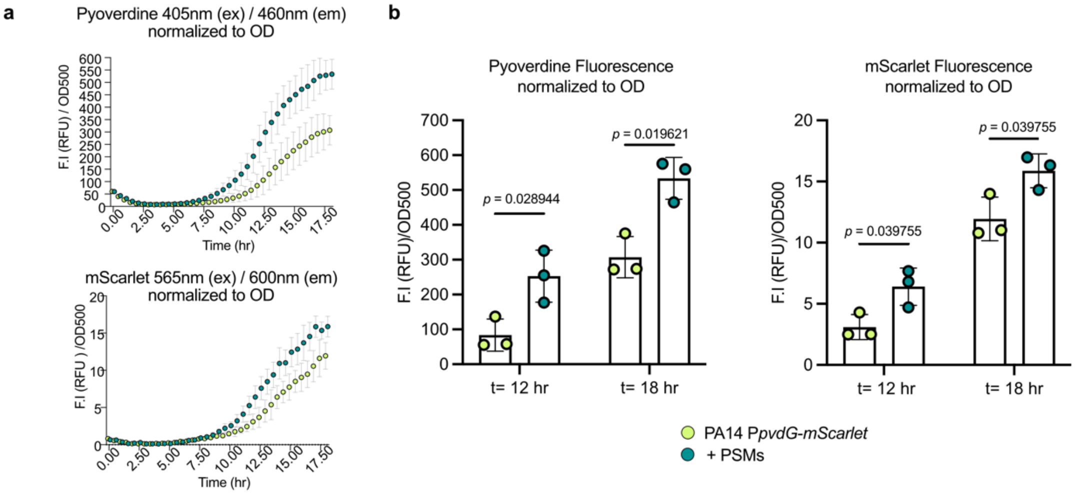


**Supplementary Figure 5. PSMs induce pyoverdine biosynthesis in *P. aeruginosa*.**

*P. aeruginosa* overnight cultures (PA14 P'*pvdG-mScarlet)* **were** diluted to O.D. (600 nm) of 0.1 in fresh M14 medium. Aliquots (100 µL) of culture were added to 96-well clear, flat-bottom polystyrene plates with or without 10 µg/ml PSMs (5 µg/ml PSMα1 and PSMα3 each), followed by incubation at 37°C with continuous orbital shaking at 1200 rpm using a microplate reader (VarioScan). Fluorescence intensity (mScarlet: 565 nm ex/ 600 nm em; pyoverdine: 405 nm ex and 460 nm em) and optical density (UV absorption at 500 nm) measurements were taken every 30 min for 18 h. Measurements for triplicate wells were averaged, normalized to growth and plotted for raw fluorescence intensity units (**a**) with statistical quantification of end time points (**b**). Data shown represent the mean and standard deviation of at least three independent replicates. Statistical significance was determined by multiple *t*-test followed by multiple comparisons test for False Discovery Rate adjusted *P*-value.

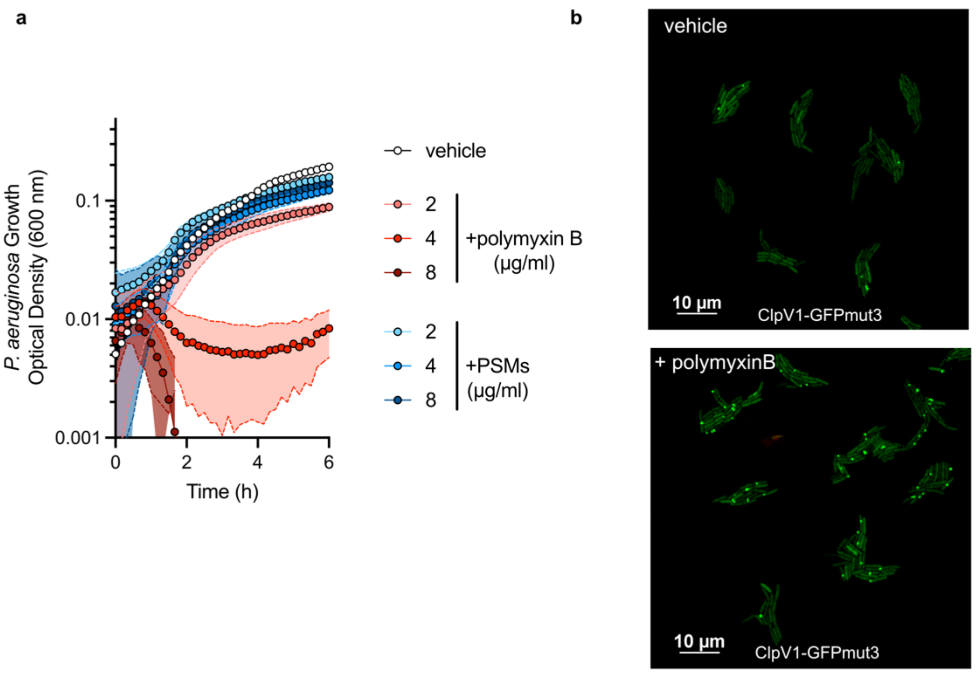


**Supplementary Figure 6. PolymyxinB inhibits *P. aeruginosa* growth and induces T6SS firing.**

**a.** *P. aeruginosa* growth over time monitored by $OD_{600}$ under indicated treatment conditions. **b.** Representative fluorescence microscopy images of *P. aeruginosa* ClpV1-GFPmut3 treated with polymyxin B.

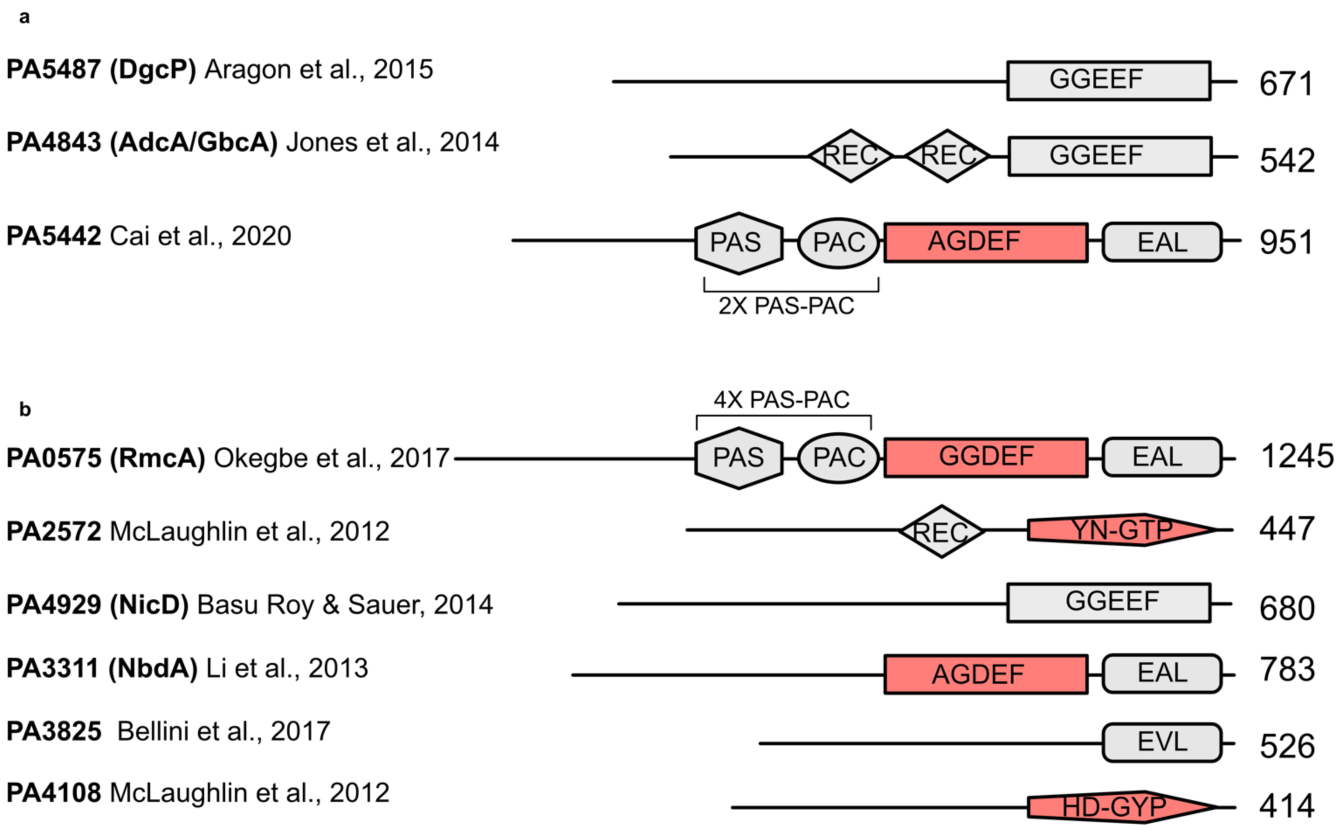


**Supplementary Figure 7. Diverse c-di-GMP sensing and regulatory enzymes[8] mediate *P. aeruginosa* interspecies response to PSMs and *S. aureus.***

Domain structures of c-di-GMP sensing/regulatory enzymes detected to be significantly up-regulated in response to PSMs pulse-in (**a**) and *S. aureus* coculture (**b**). Red color indicates silent domains. GGDEF/AGDEF/GGEEF: diguanylate cyclase (DGC) domain; EAL/EVL: phosphodiesterase (PDE) domain; PAS, PAC, REC are other sensory and/or regulatory domains frequently associated with DGC and PDE domains. For references, see Aragon et al., 2015[9], Jones et al., 2014[10], Cai et al., 2020[11], Okegbe et al., 2017[12], McLaughlin et al., 2012[13], Basu Roy & Sauer, 2014[14], Li et al., 2013[15], Bellini et al., 2017[16].

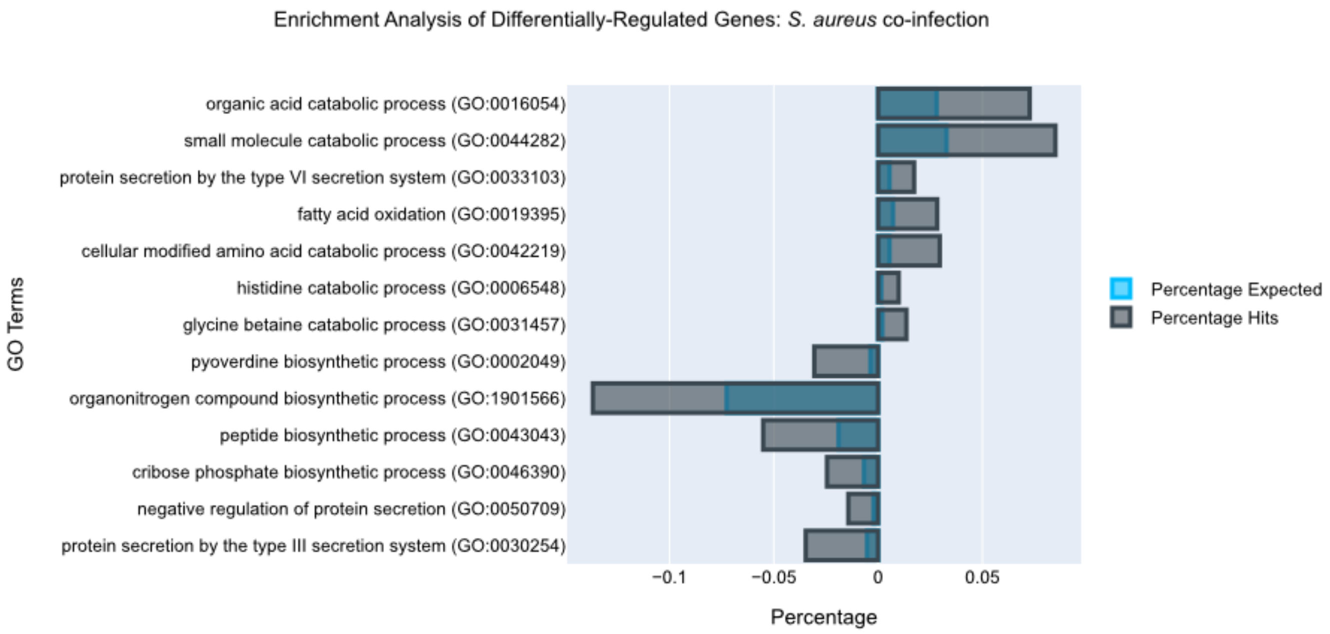


**Supplementary Figure 8. Functional enrichment analysis of *P. aeruginosa* genes differentially regulated during co-infection with *S. aureus*.**

Up-regulated (fold change > 2, *P*<0.05) and down-regulated (fold change < -2, *P* < 0.05) genes from RNA sequencing experiment were analyzed for functional classification using the PANTHER (Protein ANalysis THrough Evolutionary Relationships) Classification System (http://pantherdb.org/about.jsp). Selected functional categories that showed statistically significant enrichment (FDR adjusted P-value <0.05) are plotted. Population total (PT) refer to all protein/genes quantified in each experiment; population hits (PH) refer to all protein/genes annotated as indicated categories in each experiment; list hits (LH) refer to protein/genes annotated as indicated functional categories in subset lists; list total (LT) refer to total number of proteins/genes in each sublist. Percentage expected = PH/PT; Percentage hits = LH/LT.

Table 1. Strains and primers used in this study.

| Name | Genotype or sequence | Source | Notes |
|---|---|---|---|
| ***P. aeruginosa* strains** | | | |
| UCBPP-PA14 | WT | Laboratory collection | |
| PA14 NLL-MetRS | attTn7::mini-Tn7T-$Gm^R$ $P_{trc}$:*nll-EcmetRS* | Babin *et al*., 2017[17] | |
| PA14 *P'pvdG-mScarlet* | *P'pvdG-mScarlet* (pSB175) | Zarrella *et al*., 2022[18] | |
| PAO1 | WT | Laboratory collection | |
| PAO1 ClpV1-GFPmut3 | | Obtained from Joseph Mougous (UW) | |
| PAO1 ΔH1-T6SS | Δ*tssB1* (Δ*PA0083*) | This study | |
| PAO1 ΔH2-T6SS | Δ*hsiB2* (Δ*PA1657*) | This study | |
| PAO1 ΔH3-T6SS | Δ*hsiB3* (Δ*PA2365*) | This study | |
| ***S. aureus* strains** | | | |
| USA100 | WT | Laboratory collection | |
| USA300 LAC | WT | Laboratory collection | |
| USA300 LAC Δ*psmα*1-4 | Δ*psmα1–4* | Sayed *et al*., 2015[19] | |
| USA300 LAC Δ*psmαδ* | Δ*psmα1–4 δATG-ATT* | Sayed *et al*., 2015[19] | |
| **Primers** | | | |
| *PA0083*UpF01-attB1 | GGG GAC AAG TTT GTA CAA AAA AGC AGG CTA CGT CGC CGA ACG GGT CGG CTC | This study | For PAO1 Δ*tssB1* |
| *PA0083*UpR01 | GGA ATC CTC TTA CGC CTG CGG TCC CAT CTT GTT TCT CCC TCG CG | This study | For PAO1 Δ*tssB1* |
| *PA0083*DownF01 | CCG CAG GCG TAA GAG GAT TCC | This study | For PAO1 Δ*tssB1* |
| *PA0083*DownR01-attB2 | GGG GAC CAC TTT GTA CAA GAA AGC TGG GTA GCA GCC ATA GGG CTC GCC G | This study | For PAO1 Δ*tssB1* |

| | | | |
|---|---|---|---|
| *PA0083*F01-SEQ | CAT GCC CTG GCC ATC GAG | This study | For PAO1 Δ*tssB1* |
| *PA0083*R01-SEQ | GCG GCG ACT GGT CGA AGT | This study | For PAO1 Δ*tssB1* |
| *PA1657*UpF01-attB1 | GGG GAC AAG TTT GTA CAA AAA AGC AGG CTA CGG CCA CCC GTG GCT GGA TCA G | This study | For PAO1 Δ*tssB1* |
| *PA1657*UpR01 | GGT GGC TCA GGC GTC CTG GGA CGA GCC TTC TTT GGC CAT GGC | This study | For PAO1 Δ*tssB2* |
| *PA1657*DownF01 | TCC CAG GAC GCC TGA GCC ACC | This study | For PAO1 Δ*tssB2* |
| *PA1657*DownR01-attB2 | GGG GAC CAC TTT GTA CAA GAA AGC TGG GTA GAC CGG CTG GCC ACC GAA CTG | This study | For PAO1 Δ*tssB2* |
| *PA1657*F01-SEQ | CAC GAC GGC AGC GCA TTC | This study | For PAO1 Δ*tssB2* |
| *PA1657*R01-SEQ | CGG GCG AAC TGG GCG AC | This study | For PAO1 Δ*tssB2* |
| *PA2365*UpF01-attB1 | GGG GAC AAG TTT GTA CAA AAA AGC AGG CTA CCC AGC TCC AGG CTC CAT ACC G | This study | For PAO1 Δ*tssB3* |
| *PA2365*UpR01 | GGA AGA GGG TCA GGC CGG CTG GTG CTG CGT ACT CTC GGC CAT | This study | For PAO1 Δ*tssB3* |
| *PA2365*DownF01 | CAG CCG GCC TGA CCC TCT TCC | This study | For PAO1 Δ*tssB3* |

| | | | |
|---|---|---|---|
| *PA2365*DownR01-attB2 | GGG GAC CAC TTT GTA CAA GAA AGC TGG GTA GCA GGC TGA AGG GGT GTC CGC | This study | For PAO1 Δ*tssB3* |
| *PA2365*F01-SEQ | CCG ACT CGA TGA ACT GCC C | This study | For PAO1 Δ*tssB3* |
| *PA2365*R01-SEQ | GCG GAT GGC GAC CGA AGG | This study | For PAO1 Δ*tssB3* |
